# Thalamic regulation of a visual critical period and motor behavior

**DOI:** 10.1101/2022.09.21.508908

**Authors:** John Hageter, Jacob Starkey, Eric J Horstick

## Abstract

During the visual critical period, sensory experience refines the structure and function of visual circuits. The basis of this plasticity was long thought to be limited to cortical circuits, yet recently described thalamic ocular dominance plasticity challenges this dogma and demonstrates greater complexity underlying visual plasticity. Yet how visual experience modulates responses of thalamic neurons or how the thalamus modulates CP timing is incompletely understood. Using a novel larval zebrafish, thalamus-centric ocular dominance model, we show functional changes in the thalamus and a role of inhibitory signaling to establish critical period timing using a combination of functional imaging, optogenetics, and pharmacology. Moreover, hemisphere-specific functional changes in genetically defined thalamic neurons correlate with changes in visuomotor behavior, establishing a role of thalamic plasticity in modulating motor performance. Together, our work demonstrates that visual plasticity is more broadly conserved and shows that visual experience leads to neuron-level functional changes in the thalamus that require inhibitory signaling to establish critical period timing.

## Introduction

Early life experience is instrumental in refining brain structure and function, and discrete developmental windows known as critical periods (CP) have a prominent role in sensory-dependent plasticity. Furthermore, disruptions to sensory input or plasticity regulation can lead to impaired sensory processing and is associated with severe neurological diseases^1–3^. Ocular dominance plasticity (ODP) is a classic readout used to study visual experience-dependent plasticity^4,5^. During ODP, animals undergo monocular deprivation to produce asymmetric visual experience, which drives structural and functional changes in visual processing centers^3^. Extensive work using ODP has revealed robust structural and functional changes in binocular connectivity, synaptic scaling, and structural changes that occur between hemispheres^6–10^. ODP experiments have been performed on diverse mammalian systems, demonstrating visual plasticity across diurnal, nocturnal, and crepuscular species with varying visual acuities and circuit structures^4,11–14^. These diverse mammalian models have provided in-depth insight into the sensory-dependent changes in the primary visual cortex, illuminating distinct mechanisms incorporating excitatory and inhibitory circuits^15–19^. Conversely, subcortical regions have received significantly less attention, yet it is now known that thalamic neurons exhibit ODP in mammals^20–23^. Interestingly, the mechanisms driving subcortical ODP show distinct properties compared to visual cortex ODP, demonstrating independent mechanisms^20–23^. Subcortical contributions to sensory-driven plasticity add new layers of complexity to critical period regulation and visual plasticity, yet how subcortical neurons are modulated by early visual experience remains incompletely understood. For example, functional characterization of thalamic ODP is largely dependent on recordings from thalamic projections in layer 1 of the visual cortex or organotypic slices^21–23^. Elucidating neuron-level thalamic sensory-driven changes is challenging as in vivo analysis in deep brain regions is technically challenging and significant cortico-thalamic feedback signaling obscures resolving thalamic-specific changes. Moreover, GABAergic signaling is well-established to regulate CP onset^24^, yet thalamic contributions to CP timing are unknown.

Here we have developed a novel model for visual critical period plasticity in larval zebrafish that mirrors crucial features of mammalian ODP. Because zebrafish do not possess a cortex, the mechanisms driving subcortical plasticity can be studied independently of cortical feedback or without additional manipulations to remove feedback, providing a robust thalamic-centric ODP model. Asymmetric visual experience during a developmentally restricted time period leads to sustained neurophysiological and visuomotor change. In single animals, we show how asymmetric visual experience imposes specific visuomotor response types, which correspond with asymmetric response strengths and numbers of a genetically defined subset of thalamic neurons. Similar to well-established mammalian models^10,24^, the timing of the zebrafish critical period depends on GABAergic signaling. Our work suggests that ODP is more broadly conserved in vertebrates. Moreover, our zebrafish model recapitulates key elements associated with ODP and revealed neuron-level changes in the thalamus that instruct plasticity and behavioral performance.

## Results

### A visual critical period modulates the performance of a visuomotor asymmetry

Our previous work demonstrated that larvae zebrafish exhibit a persistent motor asymmetry following the loss of environmental illumination^25–27^. Following the loss of global visual input, an individual will preferentially utilize leftward or rightward turns (Supplementary Figure 1A). Turn bias is maintained over hours and multiple days^26^. This stereotactic behavior is consistent with search behavior patterns observed across numerous species^28–30^. In wildtype zebrafish, left and right turning types are observed in equal proportions^26,27^. Moreover, turn direction preference is also not heritable, implying stochastic molecular or environmental cues determine an individual’s turning type^26,27^. However, we previously were unable to identify a developmental ‘switch’ instructing an individual to adopt a left or right turn bias preference. Therefore, we hypothesized that asymmetric photic experience could modulate the direction of turn bias behavior. To answer this question, we temporally restricted visual input to a single eye recapitulating the ocular dominance plasticity (ODP) assay^3^. As zebrafish vision is largely monocular^31^, we controlled visual experience by embedding larvae in low melting temperature agar and positioning individuals so that either a single or both eyes were directed to a light source, we call this prep the ‘CP assay’ (Figure 1A). Larval zebrafish are well suited to this style of visual control as they acquire nutrients from a yolk and exchange oxygen and necessary ions by diffusion^32,33^. The CP assay also generated a robust visual asymmetry despite the transparency of larval zebrafish, likely due to the heavily pigmented retinal epithelium. Alignment of the eyes in the CP assay blocks approximately 58% of visible light, yielding an asymmetric visual experience (Supplemental Figure 1B-C).

**Figure 1:**
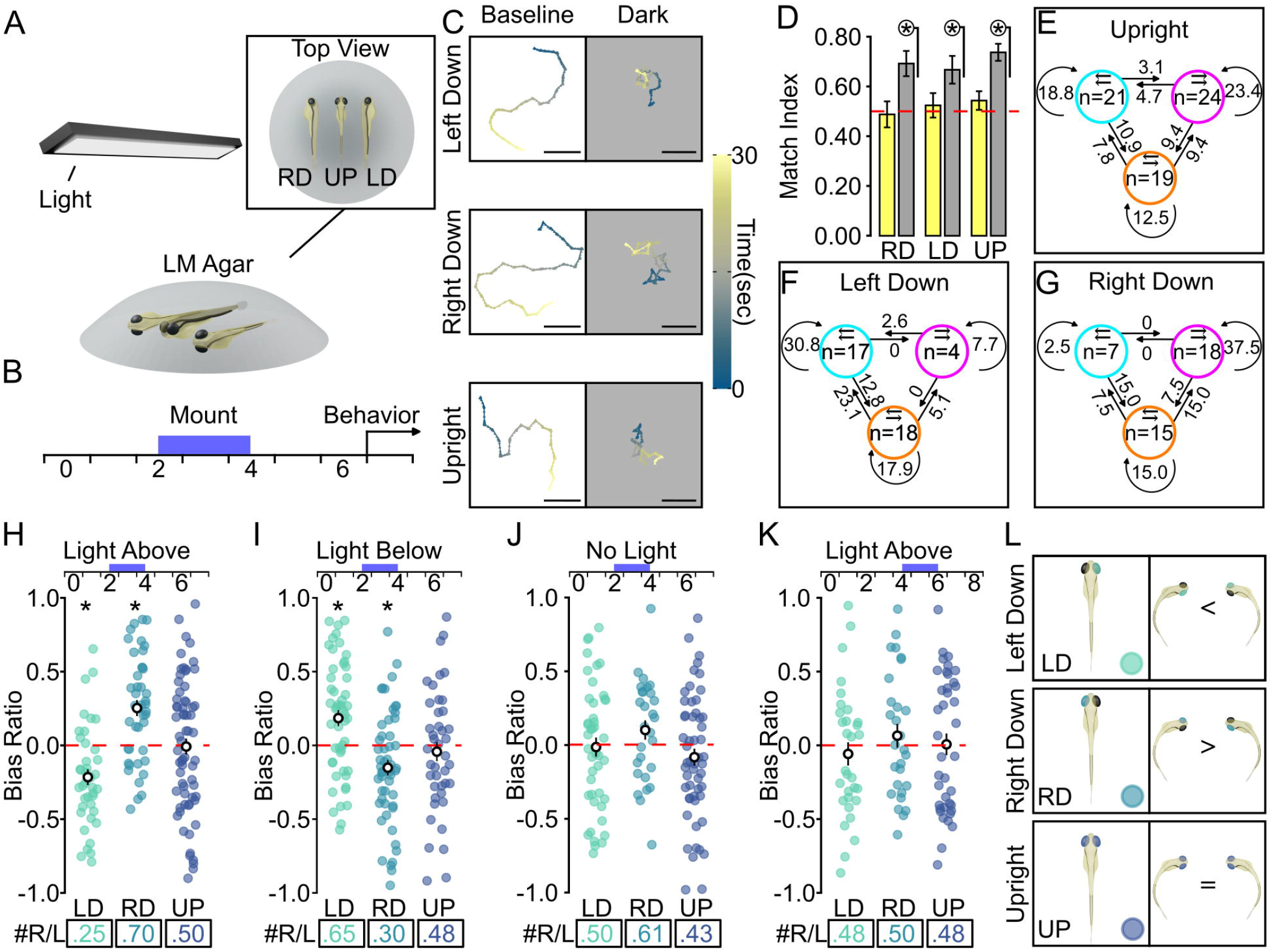
Asymmetric visual input directs larval zebrafish motor bias. **(A)** Diagram of CP assay. Inset shows an overhead view of fish and mounting orientations (RD = right side down, UP = both eyes up, LD = left side down). **(B)** Timeline of CP assay for inducing turn bias direction. Purple bar indicates the duration of embedding and visual control. **(C)** Representative tracks of mounted individuals during baseline (left) and following loss of illumination (dark, right). Scale bars=10mm. Scale bar shows time in seconds. **(D)** Match index for mounted individuals during baseline (yellow) and dark (gray) recording conditions. Dotted red line indicates random match index. Circled asterisk indicates p < 0.05 one-sample Wilcoxon signed rank test to 0.5 (RD N=40, LD N=35, UP N=62). Relative frequencies of paired directional transitions from the first two trials compared to the last two for **(E)** upright, **(F)** left-mounted, or **(G)** right-mounted fish. N’s for the first two trials are denoted in colored circles for individuals who had repeated left (cyan), right (magenta), or random (orange) trials. Paired arrows in circles indicate turn direction over the first two trials. Arrows between circles indicate transition frequency and adjacent numbers are percentages. **H-K**) Average bias ratio (averaged over four trials per individual) for individuals in the CP. Bars above show dpf and the blue bar indicates the time in CP assay. Bottom: boxes show the ratio of right-biased (+ BR) to left-biased (- BR) individuals per group. All CP assay bias ratio graphs will show these same features. **H)** CP assay from 2-4 dpf (LD=39, RD=40, UP=62) with light overhead. **I-J)** Same as H with the light source rotated below the embedded larvae in **I** (LD N=51, RD N=53, UP N=42) or with no photic experience in **J** (LD N=44, RD N=28, UP N=54). **K)** CP assay performed later in development compared to H-J. Behavior tested at 8 dpf (LD N=31, RD N=28, UP N=39). * indicated p<0.05 one-tailed t-test to 0. White circles with black vertical line show the group mean and SEM. **(L)** Diagram of visually mediated changes to turn bias direction during the CP. All error bars represent ±SEM.

To determine whether asymmetric visual experience would dictate turn bias direction, we prepared individuals in the CP assay for an approximately 48-hour window, from 2 to 4 days post-fertilization (dpf). This interval was selected as 4 dpf is the earliest turn bias is observed^27^ and coincides with retinal innervation of zebrafish visual centers^34,35^. We averaged responses over four paired light on and off recording intervals to measure turn bias and used a bias ratio (BR)^27^ to quantify the proportion of same-direction turning (Supplementary Figure 2A). During this time interval, the CP assay did not disrupt the development of turn bias behavior (Figure 1C-D, Supplementary Figure 2B). However, asymmetric visual experience during this developmental period was capable of imposing a turn bias direction (1-way ANOVA *F(2,146)=14*.*43*, p<0.0001) contralateral to the light-facing eye, whereas upright embedded larvae displayed equal proportions of left and right biased individuals (Figure 1E-H). No population-level shifts in motor asymmetry direction were observed during baseline illumination, implicating a specific effect on search motor patterns (1-way ANOVA *F(2,146)=1*.*58*, p=0.21) (Supplementary Figure 2C).

**Figure 2:**
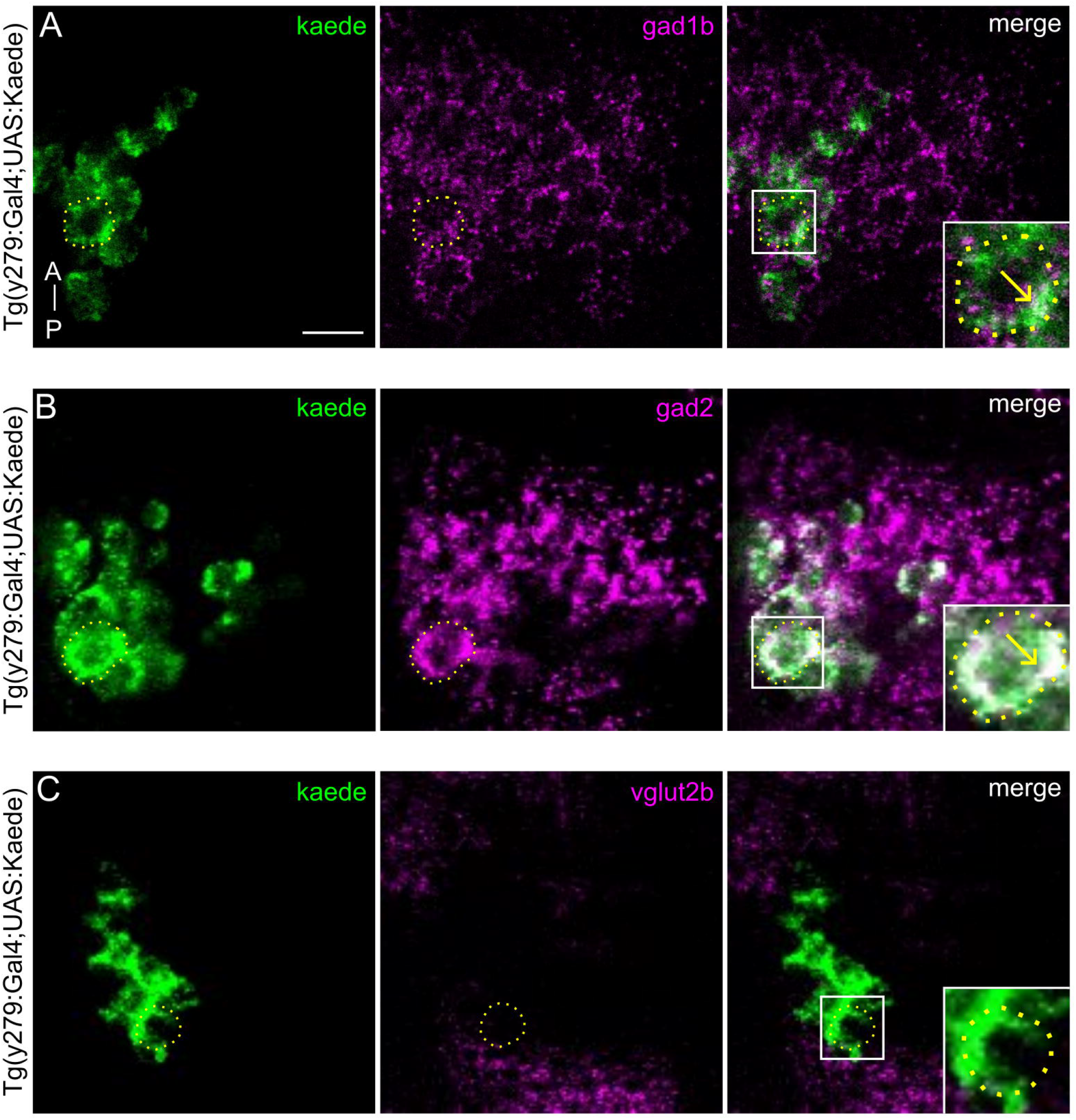
y279 specified AMNs are GABAergic. FISH HCR of *Tg(y279:Gal4; UAS:kaede)* larvae at 3 dpf. All images are single-plane confocal scans showing AMNs from a single hemisphere. **(A)** Kaede mRNA (left: green), gad1b mRNA (middle: magenta), and merged (right). Dotted circle outlines a single-cell body and arrows indicate co-localization. Similar mRNA labeling was observed across fish: gad2 (observed in N=3 larvae) **(B)** and vglut2b (observed in N=4 larvae) **(C)**. A similar pattern was observed in N=3 larvae for each label. Scale bar 10 µm.

Three key hallmarks define critical period plasticity: changes are sensory-dependent, plasticity is developmentally restricted, and change is sustained^3,36,37^. Larvae that experienced asymmetric visual input maintained a preferred turn direction for up to two weeks post-fertilization, approximately 168 hours post-initial testing (Supplementary Figure 2D-F), extending upon previously reported findings that did not control visual experience^26^, demonstrating a sustained sensory-dependent effect on motor asymmetry (turn bias direction). We next wanted to establish that the observed plasticity was photo- and not proprio-sensitive. If photic signals induce turn bias direction, we expect a correlation with light source position versus no correlation if proprioceptive signals were driving plasticity. Indeed, we observed the expected changes in motor asymmetry whether we rotated the source light in the CP assay (Figure 1I, Supplementary Figure 2G-I) or mounted larvae upright with a light source positioned to the side (Supplementary Figure 2J-M). These data show that visual experience instructs turn bias direction.

Next, we wanted to establish whether the observed plasticity was consistent with a critical period. Above, we have demonstrated that the CP assay imposes motor asymmetry direction through much of larval development. The additional key criterion for CP plasticity is that imposed changes are sensory dependent and developmentally restricted^3,36,37^. To address these other criteria, first, we prepped larvae in the CP assay in constant darkness, yet only during the embedding period. The absence of light during the critical period completely abolished turn bias modulation (1-way ANOVA *F(2,123)=1*.*86*, p=0.16), yet did not disrupt overall motor asymmetry (e.g., did not disrupt match index or total turning)(Figure 1J, Supplementary Figure 2N-P). Next, we examined if controlling visual experience at later developmental stages would impact turn bias by performing the CP assay from 4-6 dpf. Contrary to early asymmetric visual experience, an asymmetric visual experience later in development did not induce shifts in turn bias performance (1-way ANOVA *F(2,95)=0*.*57*, p=0.57) (Figure 1K, Supplementary Fig 2Q-S). Furthermore, shorter embedding intervals through 2-4 dpf were insufficient to induce turn bias direction (Supplementary Figure 3). Collectively, our experiments establish that turn bias direction during search behavior is driven by asymmetric visual experience during an early developmental critical period (Figure 1L – diagram of plasticity). Visual experience during the zebrafish visual critical period results in sustained behavioral change that is sensory-dependent and developmentally restricted – mirroring features in established ODP models.

**Figure 3:**
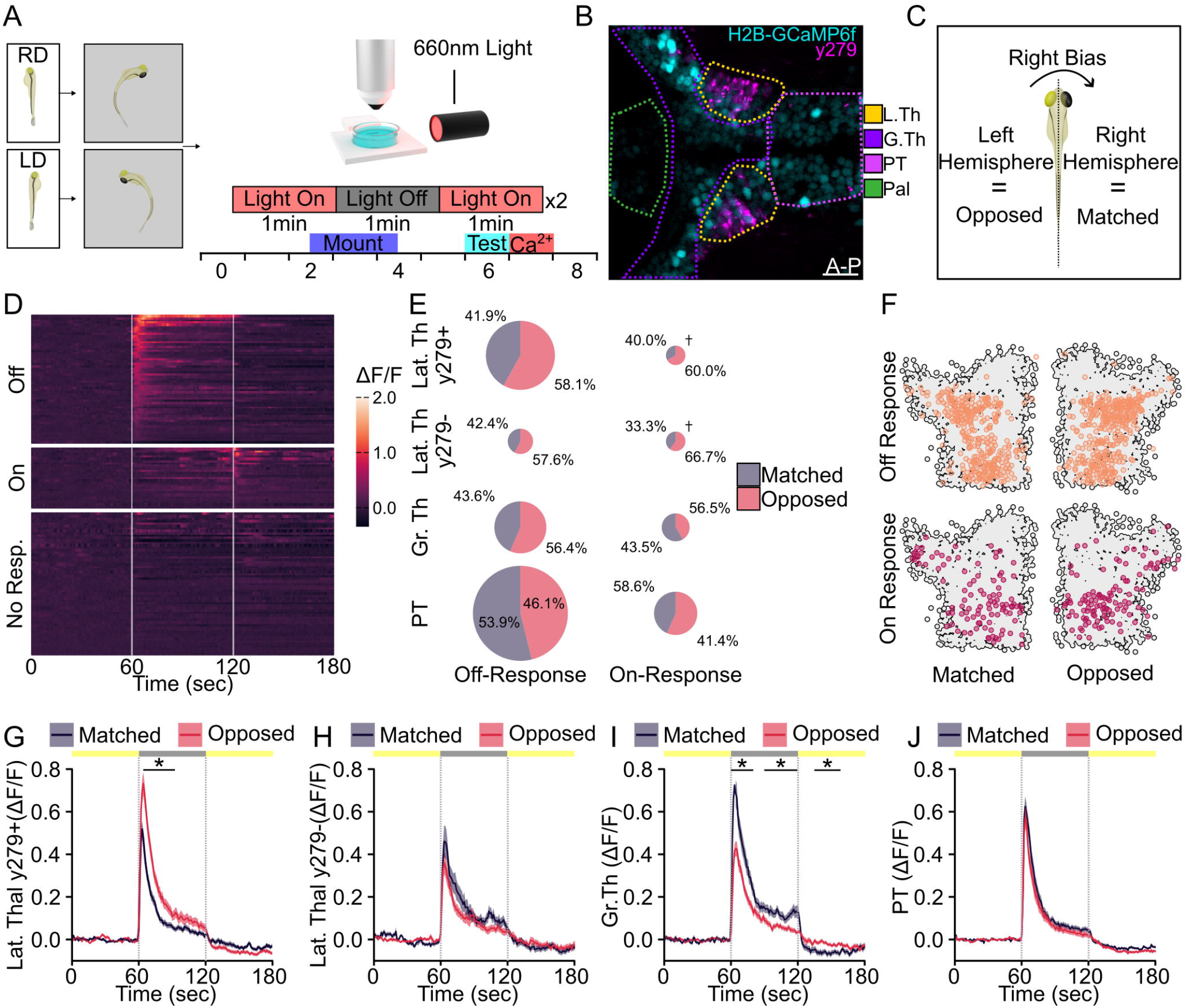
Thalamic neurons show functional asymmetries correlated to motor bias. **(A)** Workflow of panneuronal calcium imaging. **(B)** Representative segmentation of lateral thalamic (yellow), greater thalamic (purple), PT (pink), and pallium (green) neurons using *Tg(elavl3: H2B-GCaMP6f; y279 Gal4; UAS: nsfB-mCherry)*. Scale bar=20µm. **(C)** Example of matched and opposed hemispheres for a right-biased individual. **(D)** Raster plot showing responses from a subset of lateral thalamus neurons. Vertical gray bars indicate light transition (0-60 sec light ON, 61-120 sec light OFF, 121-180 sec light ON). Color scale = fluorescence change (∆F/F). **(E)** Proportions of matched and opposed visually evoked ON and OFF responses per regions. Chart sizes scaled to represent proportion of neurons represented. Cross indicates groups with fifty or fewer neurons which are set to a standard minimal scale. **(F)** Spatial distribution for matched and opposed responses between hemispheres (OFF-Response, orange N=699; ON-Response, purple N=216). Gray outline indicates the position of neurons with no light response. OFF-Response mean and standard error for **(G)** lateral thalamic y279+ (Matched N=75, Opposed N=104), **(H)** lateral thalamic y279-(Matched N=28, Opposed N=38), **(I)** greater thalamic (Matched N=61, Opposed N=79), and **(J)** PT regions (Matched N=167, Opposed N=143). Asterisk with a bar indicates a two-tailed t-test p<0.05 for at least ten consecutive time points.

### AMNs are GABAergic thalamic neurons

Our next goal was to explore the circuit-level mechanisms that support this CP. Previously, we identified approximately 60 neurons genetically labeled by the *Tg(y279:Gal4)* enhancer trap, which respond to the loss of illumination, and following bilateral ablation, causes a loss of persistent motor asymmetry, yet other search motor patterns remain intact^26^. Here we refer to these neurons as asymmetry-maintaining neurons (AMNs). As AMNs underlie the maintenance of motor asymmetry, these neurons are ideal candidates for modulating behavior in a sensory-dependent manner. Initially, we postulated AMNs reside in the anterior lobe of the posterior tuberculum based on available zebrafish brain atlas resources^26,27^. However, based on a combination of genetic and circuit structure evidence, we now propose these neurons reside in the anterior thalamus. Recent studies in larval zebrafish have demonstrated that the s1020t Gal4 enhancer trap labels the anterior thalamus, which is positioned immediately posterior to the zona limitans intrathalamica (ZLI)^38^. In addition, subsets of zebrafish thalamic neurons project and transmit illuminance information to the habenula and tectum^39,40^. AMNs are also positioned posterior to the ZLI, and using a current zebrafish single-neuron atlas and neuroanatomy tools, we show AMNs overlap with the 1020t Gal4 driver and neurons within this region project to the habenula and tectum^39,40^ (Supplementary Figure 4A-D). Our previous experiments show that a subset of y279-positive AMNs extend projections to the habenula, consistent with other reports describing thalamic projections from this region^26^. Last, the teleost anterior thalamus is composed predominately of GABAergic neurons^38^. Using HCR probes against GAD1B, GAD2, and vGlut2b, we identified that AMNs and surrounding neurons are GABAergic (Figure 2, Supplemental Figure 4E-F). Therefore, based on several lines of evidence, the y279-positive AMNs are anterior thalamus neurons.

**Figure 4:**
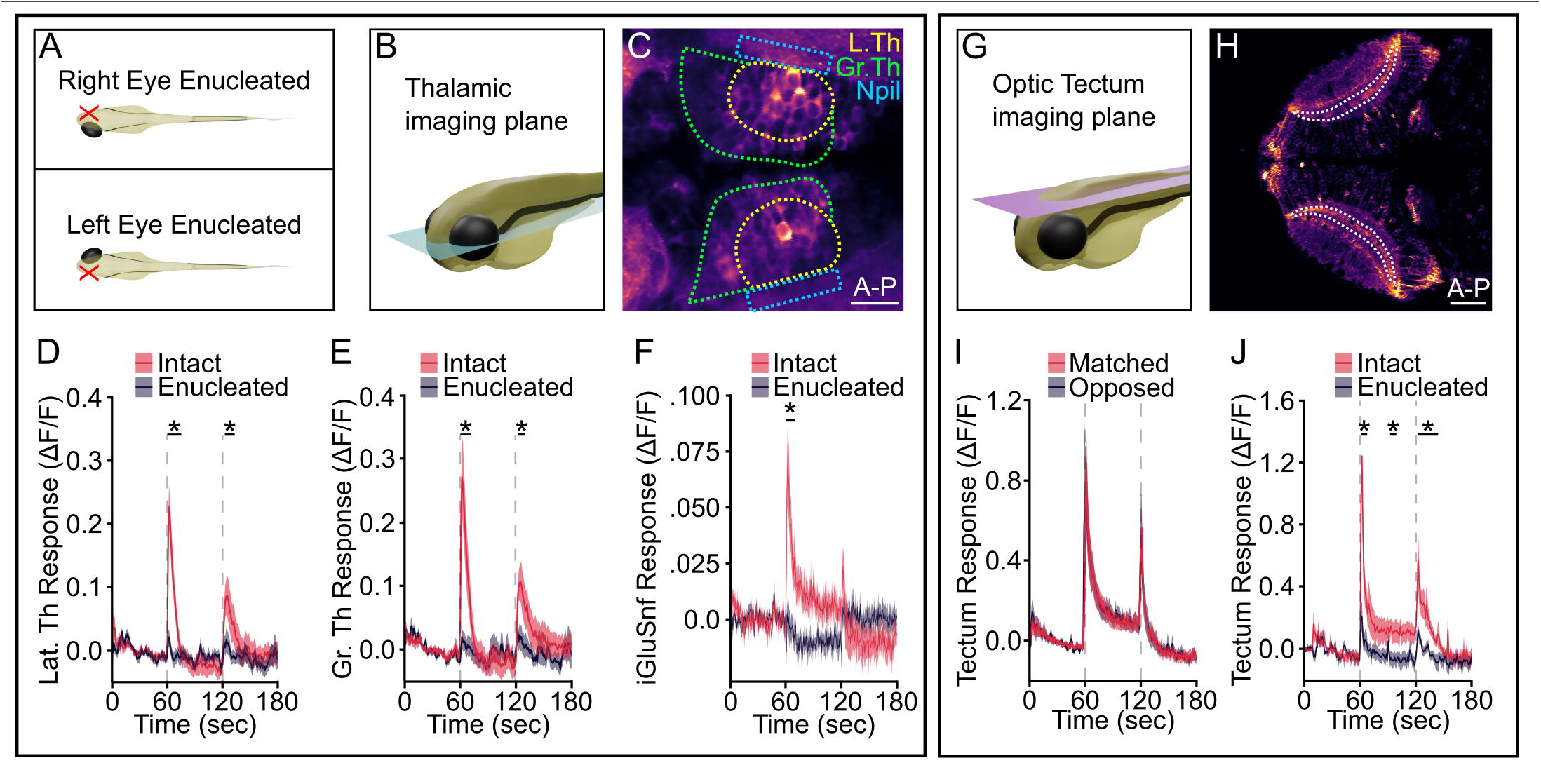
AMNs respond to visual input. **(A)** Schematic for unilateral enucleation. **(B)** Representative showing plane for thalamic neuron imaging. **(C)** Illustrative example showing regions for analysis in D-F (lateral thalamus (L. Th, white), greater thalamus (Gr. Th, green) and AMN neuropil (Npil, orange)). Scale bar=20µm. Standardized calcium responses to visual stimuli in unilaterally enucleated larvae. Vertical dotted lines lights denote the time of light OFF (60 sec) and light restoration (120 sec). Solid lines are an average and envelop shows ±SEM. **(D)** Calcium responses in the lateral thalamus (N=11) in the intact (red) and enucleated (grey) hemisphere. **(E)** Same as C showing greater thalamus (N=11) responses. **(F)** iGluSnfer responses from intact (red) and enucleated (grey) hemispheres recorded from the adjacent AMN neuropil (N=9). **(G)** Representative showing plane for optic tectum imaging. **(H)** Illustrative region used for recording calcium responses in the optic tectum neuropil. Scale bar=50µm. **(I)** Neuropil responses for hemispheres matched to the direction of turn bias (red) or opposed (purple) (N=13). **(J)** Neuropil responses contralateral to enucleated eye (red) or ipsilateral (purple) (N=11). Asterisk and bar, two-tailed t-test p<0.05 for at least three consecutive time points.

### Thalamic asymmetries correlate to motor asymmetry

Since a persistent motor asymmetry is observed, we posited that an underlying functional asymmetry is required to support this behavior. Our prior studies established that y279-positive AMNs maintain motor asymmetry and respond to changes in illumination, yet no correlated asymmetries in neuron function were found that would suggest a direct instructive role^26^. In these earlier studies, a mosaic labeling strategy was used, which may have missed a behavioral correlation. Therefore, we performed pan-neuronal calcium imaging on larvae following the CP assay to identify if AMN activity correlated with motor asymmetry. For recordings, we employed two-photon imaging and used a red LED to elicit light-evoked neural responses, which we previously demonstrated allows the recording of light-evoked neural responses^26^ (Figure 3A). For calcium imaging, we used the transgenic line *Tg(H2B-GCaMP6f)* to resolve single-neuron responses^41,42^. We crossed this line into y279:Gal4; UAS:*NfsB*-mCherry (used as an AMN marker) to identify AMNs from surrounding thalamic and non-thalamic neurons. For analysis, single neuron responses were grouped into four regions based on neuroanatomical markers^43^: lateral thalamus (includes AMNs), greater thalamus, pallium, and posterior tuberculum (Figure 3B-C). The lateral thalamus encompasses the AMNs (y279^+^ and y279^-^), whereas we use the term greater thalamus to refer to the remainder of thalamic neurons medial and anterior of the AMNs. We recorded from a total of 6233 neurons and detected photo-mediated changes in neural activity across imaged regions (Figure 3D). Across most recorded regions, we detected three different photo-mediated response profiles (OFF-responses, ON-responses, and OFF-ON responses) that match established RGC response types^35,44^. Predominately OFF-responses were observed across tested brain regions, except for the pallium, which had minimal photo-driven outputs (Supplementary Figure 5A). To determine how photo-responsive neurons may establish motor asymmetry, we compared the numbers and positions of light-responsive ON or OFF neurons in the matched or opposed hemisphere relative to the turn bias direction (see Figure 3C). Only y279-specified AMN OFF-responders showed a hemispheric difference in neuron numbers (χ^2^(1) = 4.698, p = 0.0302), with approximately 14% more neurons in the hemisphere opposed to turn bias direction (Supplementary Table 1)(Figure 3E-F). This data shows that light OFF-responsive neurons are a predominant thalamic neuron population with an asymmetric distribution of y279-positive neurons in the lateral thalamus.

**Figure 5.**
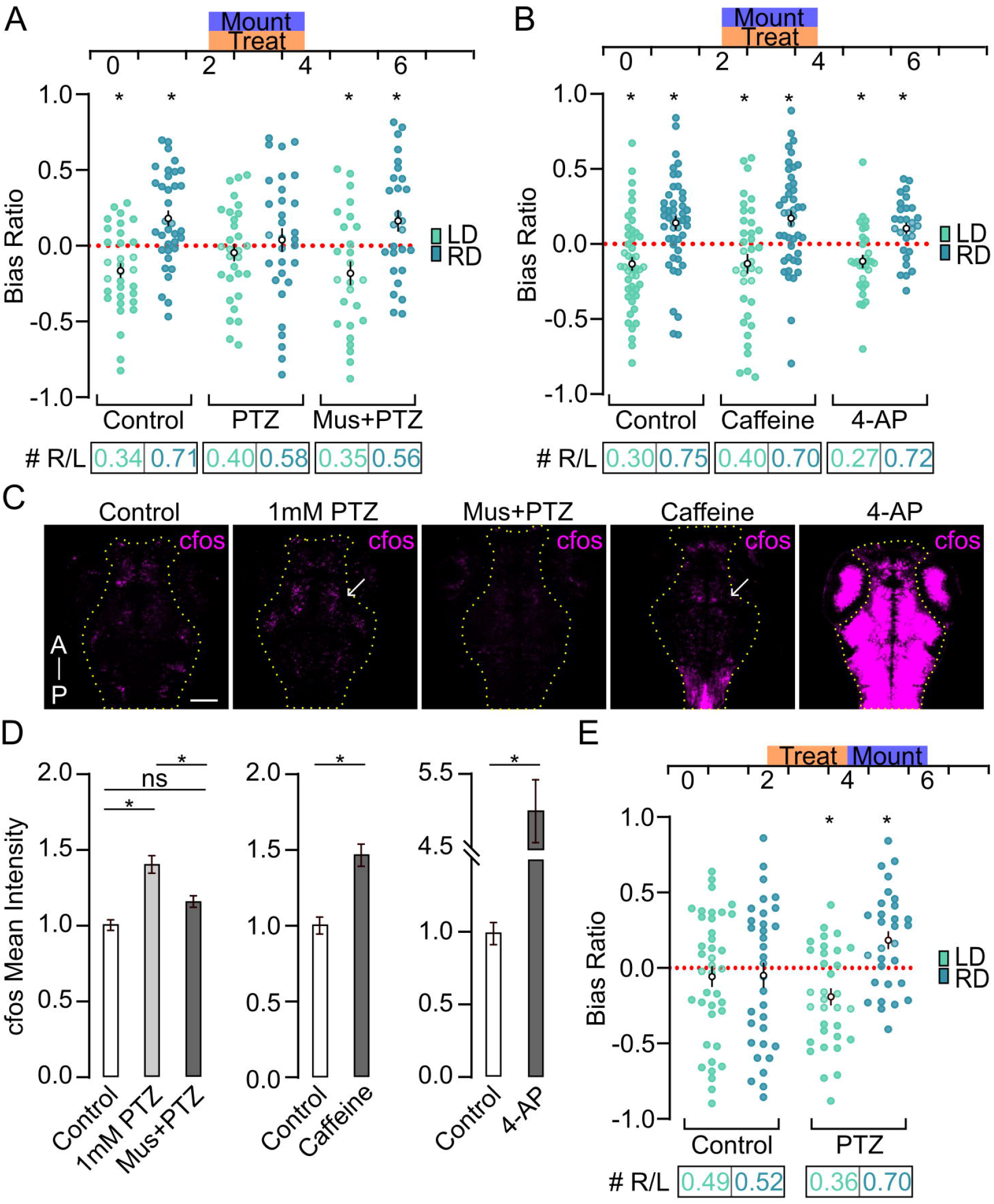
GABAergic signaling controls critical period timing. **(A)** Average BR of larvae following asymmetric visual experience. Schematic (top) shows the time in dpf of CP assay mounting (purple) and drug treatments (orange). Control (LD N=32, RD N=35), PTZ (LD N=30, RD N=31), and Muscimol + PTZ (LD N=26, RD N=27). Bottom R/L values show the proportion of left BR (negative values) to right BR (positive value) individuals. Red line indicates random movement. **(B)** Same as in A with different activity-modulating drug treatments (control LD N=46, RD N=47; 500 µM Caffeine LD N=37, RD N=43; or 400µM 4-AP LD N=30, RD N=29). **(C)** Representative c-fos mRNA labeling following drug treatments. Single plane confocal scans of 4 dpf larvae. Arrow indicates areas of increased c-fos expression. Yellow outline shows the area analyzed for c-fos signal quantification. Scale bar 100 µM. **(D)** Quantification of C. Average mean intensity of c-fos from untreated controls (N=41), PTZ (N=16), Muscimol + PTZ (N=14), Caffeine (N=14), and 4-AP (N=12). Values standardized to control. **(E)** Same as in A with larvae prepped in the CP assay from 4-6 dpf (see timeline above) (control LD N=39, RD N=33; PTZ LD N=33, RD N=30). Asterisk p<0.05 to 0.0. Asterisk with lines in D shows p<0.05 between groups.

Next, we examined whether changes in neuronal response strength paired with turn bias direction. Within the lateral thalamus, 15.2% (179/1174) of y279-positive AMNs showed an OFF-response, which coincides with the onset of motor asymmetry. Interestingly, when grouped based on behavioral performance, we find that y279-positive AMNs show a stronger response in the hemisphere opposed to the turn bias direction (Figure 3G). This analysis was based on a single neuron comparison across all tested fish; similar trends were observed when we analyzed y279-positive AMN responses on a per-fish basis (all responses averaged per larva) or per behavior (responses from left and right-biased individuals) (Supplementary Figure 5B-D). To independently confirm the observed AMN responses observed in the pan-neuronal imaging, we injected *Tg(y279:Gal4)* larvae with a UAS:H2B-GCaMP6s construct to mosaically and unambiguously label y279-positive AMNs. Using an identical recording strategy as above, the mosaic labeling of AMNs recapitulated the pan-neuronal imaging, confirming the correlation between AMN physiology and turn bias (Supplementary Figure 5E). Conversely, OFF-responses in lateral thalamic y279 negative neurons showed no hemispheric asymmetries associated with behavior (Figure 3H).

Last, we examined whether the functional asymmetries observed in the y279-positive AMNs were specific to the lateral thalamus or a broader feature in the brain. Surprisingly, OFF-responses in the greater thalamus (6.1% (140/2281) of neurons) also showed functional asymmetries that correlated with turn bias direction, yet displayed increased responsiveness in the hemisphere matched to turn bias, opposite the pattern observed in lateral thalamic y279-positive AMNs (Figure 3I). Contrary to thalamic neurons, OFF-response posterior tuberculum neurons (posterior to the thalamus; 310/2073, 15.0% of neurons) demonstrated no hemispheric asymmetries based on turn bias direction (Figure 3J). We did not explore pallium functional asymmetries as so few neurons responded to visual stimuli. Our data establish that distinct subsets of thalamic neurons are functionally correlated with motor asymmetry. Moreover, this functional asymmetry is likely a unique feature of the thalamus as surrounding brain regions show limited or uncorrelated responses with turn bias onset or direction. Together, our data suggest that visual experience changes the strength and number of OFF-responsive neurons in the thalamus.

### Thalamic AMN responses are retina dependent

Previously, we showed retinal ganglia cells (RGC), and AMN neurites colocalize in the adjacent neuropil, yet whether the retina drives AMN activity was not determined^26^. Based on mammalian ODP models and our behavioral assays in zebrafish, we hypothesize that lateral thalamus photo-responsiveness is retina driven. To test this hypothesis, we perform unilateral enucleations to eliminate visual input from a single eye and simultaneously record thalamic responses to visual stimuli in both hemispheres (Figure 4A-C). In zebrafish, RGCs project exclusively to the contralateral hemisphere^31,45^; therefore, lateral enucleation eliminates contralateral photic input. Contralateral hemisphere (enucleated eye) responses in the lateral and greater thalamic responses are lost yet maintained in the ipsilateral hemisphere (Figure 4D-E). Therefore, our data show that thalamic responses are dependent on retinal input. As RGCs are predominately glutamatergic, we next expressed the glutamate sensor iGluSnfer^46^ in AMNs and recorded from the adjacent neuropil in unilaterally enucleated larvae (Figure 4C, F). Indeed, we find glutamatergic activity following the loss of light from the intact yet not enucleated neuropil (Figure 4F). As a control, we quantified iGluSnfer responses in a posterior neuropil position away from potential RGC inputs. Away from the RGC containing neuropil no evidence of hemispheric differences in glutamatergic signaling was observed (Supplementary Figure 5E). These experiments suggest AMNs are receiving visual OFF signals from RGCs.

A potential caveat is that AMN asymmetries are driven by asymmetric retinal output created by the imbalanced visual experience during the CP assay. Because we observe different patterns of functional asymmetry across the thalamus (see Figure 3), we consider this outcome unlikely. Nevertheless, we recorded activity in the optic tectum following changes in illumination (Figure 4G-H). The optic tectum is the primary visual processing center in zebrafish and receives the majority of RGC input^47^. Unlike AMNs or the greater thalamus, optic tectum responses across hemispheres show no differences based on turn bias direction (e.g., symmetric across hemispheres) (Figure 4I-J). This data indicates that thalamic responses depend on the retina, yet sensory-driven changes in thalamus responses are not a product of asymmetric retinal output.

### GABAergic signaling maintains zebrafish visual critical period timing

Shifts in the developmental window for critical period plasticity can be induced by pharmacologic manipulation of GABAergic activity^10,18,24^, showing that the timing of GABAergic circuit development is a fundamental mechanism for regulating ODP. A recent *Drosophila* study also identified a critical period for motor behavior, which similarly requires inhibitory signaling^48^, suggesting inhibitory signaling is an essential and conserved mechanism in critical period regulation, regardless of cortical input. To test if GABAergic signaling is necessary for zebrafish ODP, we used pentylenetetrazol (PTZ), a reversible non-competitive antagonist of GABA_A_ receptors^49^. In zebrafish, high concentrations of PTZ induce seizures, resulting in clonus-like convulsions and epileptiform brain discharges^50,51^. To treat zebrafish during the entire critical period and avoid severe dysregulation of neural activity, we used 1mM PTZ, which is below concentrations reported to induce seizure^50^, and treatments maintain grossly normal motor behavior and motor asymmetry (Supplementary Figure 6A-C). To confirm and quantify activity changes, we developed a fluorescent in-situ hybridization probe to detect changes in c-fos expression, a well-establish marker for increased activity in zebrafish^52–55^. Using a high dose of PTZ, we confirmed that our HCR c-fos probe would detect increased neural activity (Supplementary figure 6D-E).

**Figure 6:**
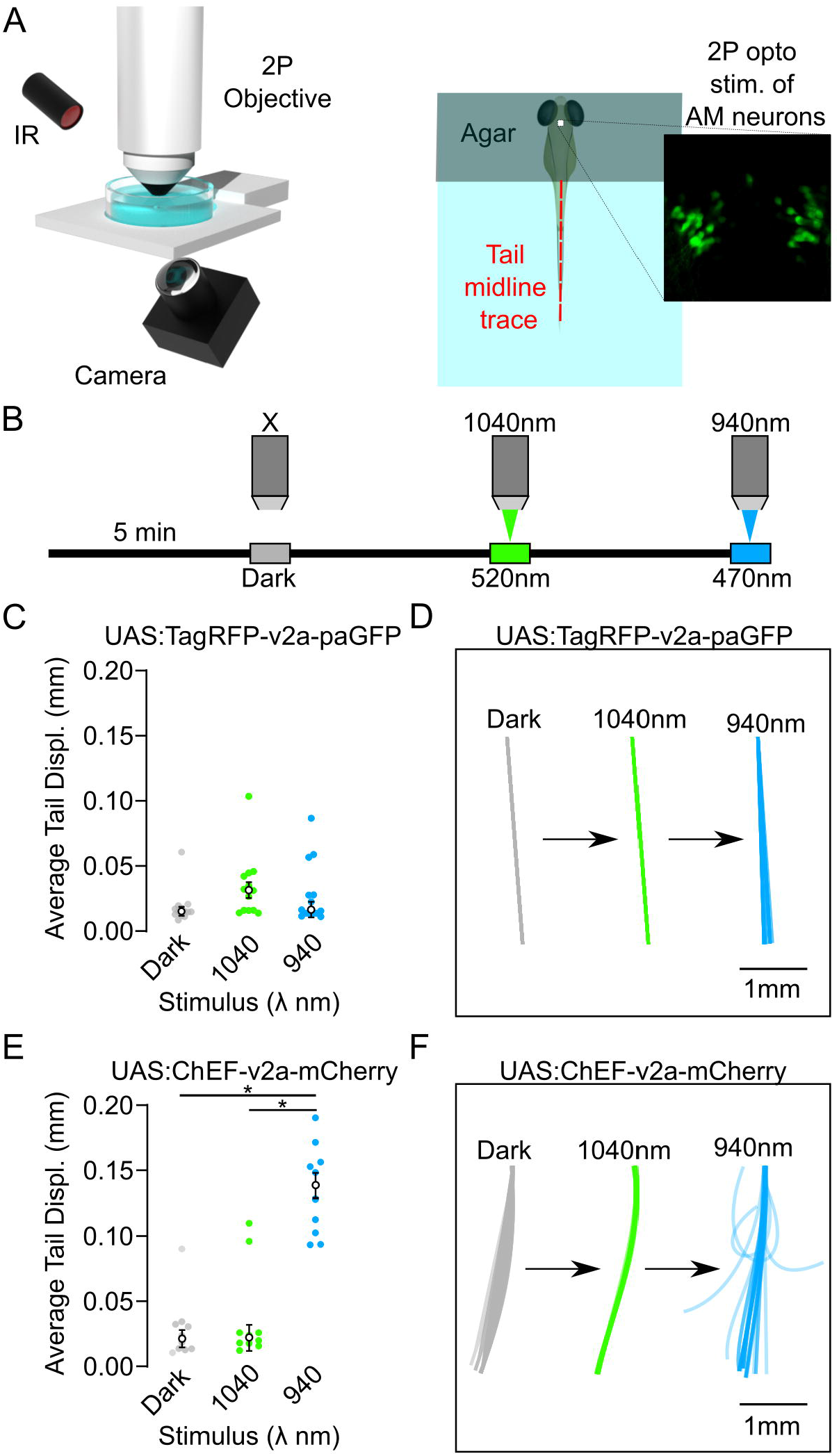
AMN activation drives motor responses. **(A)** Schematic of two-photon setup for optogenetic stimulation and tail recording (left). Diagram of head-embedded larvae and field of view containing AMNs (right). **(B)** Photic stimulation series and wavelengths. **(C)** Average tail displacement (mm) of UAS:TagRFP-v2a-paGFP (control) larvae following no stimulation dark controls (gray), 1040nm (green), and 940nm laser (blue) exposure. Individuals tested N=13. **(D)** Representative of the tail traces for analysis in C) during dark control, 1040nm, and 940nm recording. **(E-F)** Same as above for UAS:ChEF-v2a-mCherry injected individuals (N=10). Asterisk with lines shows p<0.05 between groups.

Low-dose PTZ treatment during the zebrafish visual critical period blocks sensory-dependent modulation – resulting in no population level induction of turn bias direction (1-sample t-test to 0, RD: *t(30)=0*.*50*, p=0.62; LD: *t(29)=0*.*86*, p=0.40) (Figure 5A, Supplemental Figure 7A-C). Cotreatment with muscimol, a non-competitive GABA_A_ agonist^56,57^, restores plasticity suggesting a GABA specific effect (1-sample t-test to 0, RD: *t(26)=0*.*37*, p=0.04; LD: *t(25)=2*.*34*, p=0.03) (Figure 5A, Supplementary Figure 7D-F). Interestingly, this rescue was not observed using gaboxadol, an extra-synaptic GABAergic agonist^58–60^ (Supplementary Figure 7G-I). To confirm that the loss of plasticity is GABA-specific and not a by-product of altered neural activity, we treated larvae during the CP assay with 4-AP (voltage-gated potassium channel blocker) or caffeine (adrenergic receptor agonist) to stimulate neural activity^61,62^, which we confirm using HCR c-fos labeling (Figure 5B-D, Supplementary Figure 7J-L). Despite increases in neural activity, neither 4-AP nor caffeine blocked turn bias induction following asymmetric visual experience (Figure 5B). These results support our prior experiments showing GABA_A_-specific regulation of zebrafish visual plasticity. In mammals, disruption of GABAergic signaling delays CP onset^24^, yet whether this mechanism is specific to the cortex and/or thalamus is unknown. Therefore, using our thalamus-centric zebrafish model, we next asked whether GABAergic inhibition during the established abolished plasticity or caused a delay in CP onset. To determine how GABAergic signaling effected the critical period, we PTZ treated free-swimming larvae during the established critical period (2-4 dpf), removed drug, and performed a delayed embedding in the CP assay from 4-6 dpf. Untreated control larvae showed no plasticity following 4-6 dpf CP assay, consistent with prior experiments (see Figure 1K). Conversely, following GABAergic inhibition, the period for sensory-dependent plasticity was shifted, resulting in visual plasticity at later developmental stages (Figure 5E). Our data show that 1) GABAergic signaling is necessary to establish critical period timing in zebrafish, 2) synaptic GABAergic receptors are specifically required, and 3) the role of inhibitory signaling is likely conserved across vertebrates and invertebrates. Moreover, our data implicates that thalamic inhibitory signaling contributes to CP timing.

**Figure 7:**
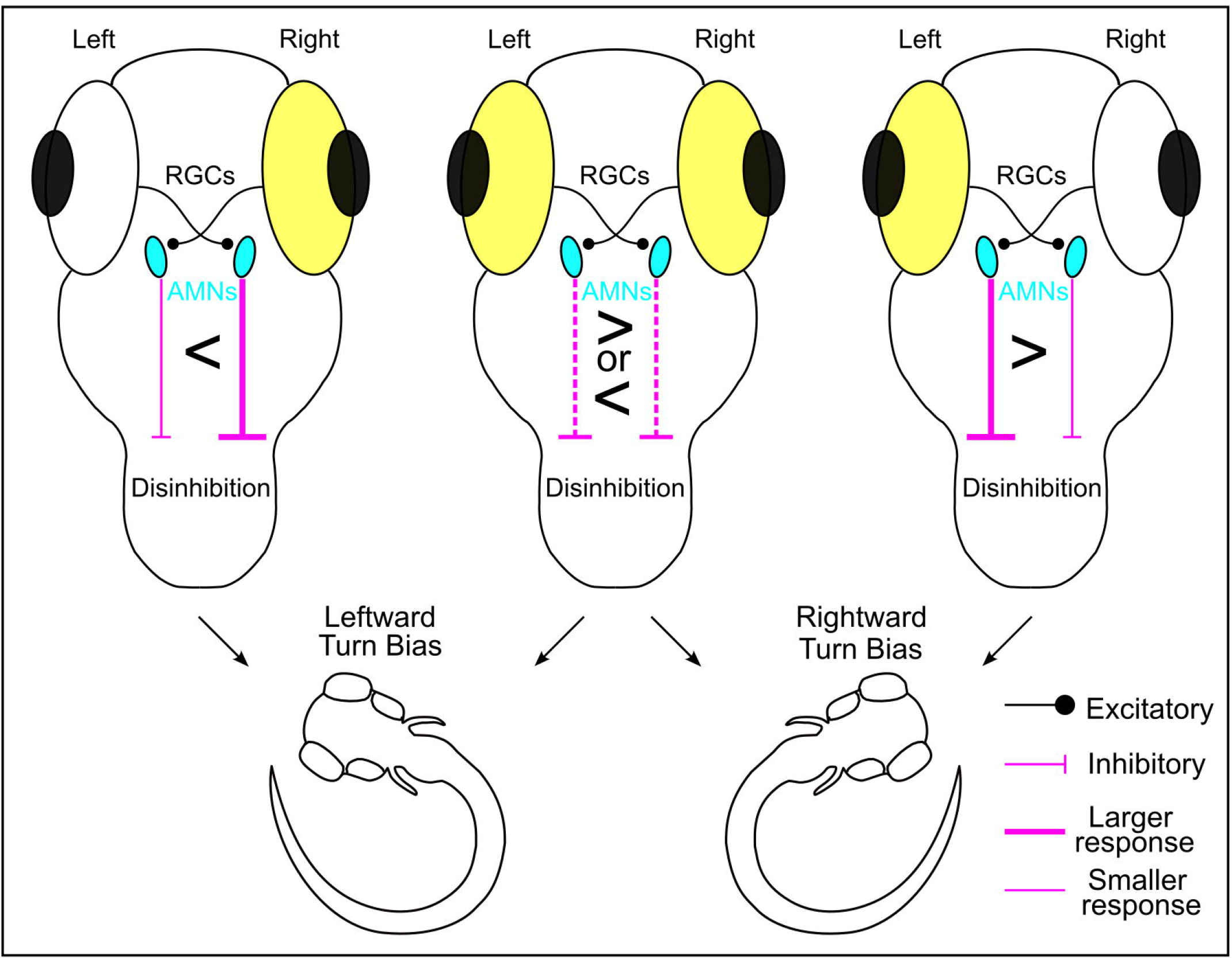
Circuit diagram. Representative diagram of proposed circuit model and visual experience impact on turn bias. Excitatory connections are shown with circled ends and inhibitory connections with flat line ends. Examples of visual plasticity are shown for right-eyed stimulated, both eyes stimulated during upright controls (middle), and left-eye stimulated. Stimulated eye shown in yellow. GABAergic AMNs from the occluded eye have increased activity compared to the visually stimulated eye. Line thickness displays response strength. We propose asymmetric descending AMNs disinhibitory signaling to premotor neurons releases motor activity to establish turn bias direction.

### Activation of AMNs sufficient to drive motor responses

Next, we wanted to determine how thalamic AMNs affect motor behavior. We focused on AMNs as this population had more OFF-responsive neurons and differences in neuron numbers between hemispheres (see Figure 3, Supplementary Table 1). Based on our functional imaging, we hypothesized that AMNs could modulate motor performance through two possible mechanisms: 1) asymmetric inhibition or 2) asymmetric disinhibition. We consider the first mechanism unlikely. Even though the stronger GABAergic AMN responses are observed in the hemisphere opposed to turn bias direction, supporting an asymmetric inhibition model for the observed turn biases, we previously demonstrated that unilateral ablation of AMNs imposes a turn bias toward the intact hemisphere (away from the ablated hemisphere)^26^. Therefore, if AMNs controlled motor behavior through asymmetric inhibition, a turn bias toward the ablated side would be expected and was not observed^26^. Therefore, we used optogenetic tools to directly manipulate AMN activity to address whether AMNs modulate motor performance using disinhibition. We injected y279 larvae with a UAS:chEF-v2a-mCherry driver for optical stimulation^63^, UAS:TagRFP-v2a-paGFP (used only for TagRFP expression) as a control, or UAS:GtACR2-tdTomato for optical inhibition^64^. Because the y279 line labels neurons in addition to AMNs^26,27^(see Supplementary Figure 3), we used a two-photon laser to photo-stimulate AMNs with three dimension precision while simultaneously recording tail movement as a readout of motor responsiveness (Figure 6A). We have shown both brain hemispheres can support motor asymmetry using unilateral ablation^26^, and expression of the optogenetic drivers is mosaic, so we bilaterally stimulated AMNs because interpreting hemisphere-specific stimulation would be challenging. Because two-photon lasers are tunable to specific wavelengths, we performed a stimulation series on each larva, including a baseline (no laser), stim-ʎ off-peak (1040nm 2P/520nm at cells), and stim-ʎ on peak (940nm 2P/470nm at cells) (Figure 6B). If AMNs modulate motor behavior using disinhibition, we expect direct activation of AMNs to release motor activity (tail movement) in our preps. Indeed, larvae expressing the optogenetic activator chEF, exposing AMN neurons to stim-ʎ on peak (940nm) evoked tail movement, which was not observed during baseline or off-peak laser exposure (1-way ANOVA *F(2)=54*.*93*, p<0.0001)(Figure 6C-D). Across all photo-stimulation intervals, no significant tail movement was observed in controls or larvae expressing the optogenetic inhibitor GtARC2 (Figure 6E-F, Supplementary Figure 8A-B). We confirmed our two-photon stimulation paradigm did not result in neuronal death (Supplementary Figure 8C-D), suggesting motor changes were due to optogenetic stimulation. These data show that thalamic AMNs can affect motor activity by disinhibiting downstream motor control pathways.

## Discussion

We demonstrate a novel visual critical period in larvae zebrafish, providing a thalamus-centric model to characterize visual plasticity mechanisms. Our zebrafish model recapitulated hallmarks of mammalian ODP such as using asymmetric visual experience, sensory-dependent regulation, sustained functional changes, GABA signaling dependent, and developmentally restricted. Notably, asymmetric visual experience in zebrafish imposes physiological asymmetries in the thalamus, providing a first neuron-level functional analysis of ODP-driven changes in thalamus neurons to the best of our knowledge. In mammals, the thalamus receives significant cortico-thalamic feedback, and isolating thalamic changes requires extensive manipulation^21,22^. Conversely, zebrafish lack a cortex^65^, eliminating cortico-feedback. Therefore, zebrafish is a robust model for establishing foundational subcortical plasticity principles.

In zebrafish, experience-dependent changes in the nervous system are well documented. For example, in the optic tectum, environmental light is necessary to shape tectal activity patterns^66,67^. One study suggests a 5-7 dpf critical period for tectum functional development^66^, yet does not examine asymmetric visual experience. Conversely, asymmetric visual experience has been investigated in *Xenopus* tadpoles, which drives changes in isthmic-tectal communication^68^, in which the tectum is the homolog of the superior colliculus^69^. In mammals, minimal ODP has been described in the superior colliculus^70^. In larval zebrafish, we did not observe functional asymmetries in the optic tectum. Nor did we observe functional asymmetries outside the zebrafish thalamus, suggesting that asymmetric visual experience leads to changes only in the thalamus. In zebrafish, the thalamus controls various photo-mediated responses^39,40^. Because only a subset of y279-positive thalamic neurons responded to the light-OFF stimulus, neurons across the thalamus are likely tuned to specific photic cues. Indeed, a subset of thalamus neurons project to the tectum and regulate loom-induced startle in zebrafish^40^, and in mammals, portions of the thalamus respond to illuminance information to control defensive behavior^71^. In a previous study, we did not observe AMNs extending projections to the tectum^26^, supporting the hypothesis that neurons in the zebrafish thalamus are divided into unique photo-responsive subsets and suggesting AMNs operate in a tectum-independent manner, consistent with the lack of turn bias correlated functional asymmetry in the zebrafish tectum (see Figure 4).

### Comparison to mammalian plasticity

Despite the benefits of the zebrafish as a model, there are differences when compared to mammalian ODP. For example, changes in binocularity contribute to the functional shifts observed in mammalian visual processing centers following asymmetric visual input as both hemispheres receive contralateral and ipsilateral RGC connections^4,19,37,72,73^. Conversely, zebrafish RGCs project to the contralateral hemisphere, providing only monocular input^31,74^. However, RGC input to the mammalian thalamus is predominately monocular^21,23^. Therefore, pertinent ODP changes in zebrafish are also likely occurring in the thalamus despite differences in binocularity. The neuronal composition also differs between species. In mammals, excitatory relay neurons in the dorsal lateral geniculate nucleus (dLGN) are the primary thalamic retinorecipients that project to the visual cortex, and these projections within the cortex have been the primary target for functionally characterizing thalamic ODP^21,23^. A subset of inhibitory dLGN interneurons also receives RGC input^37,73^, yet ODP changes have not been tested. Conversely, the zebrafish thalamus is predominately GABAergic yet is also a retinorecipient target^38,47^. Our previous and current work suggests AMNs likely receive RGC input. Therefore, in zebrafish, ODP changes are likely focused on GABAergic neurons due to the lack of a cortex and putative need for a relay neuron homolog. In zebrafish, a clear LGN homolog has not been identified^38^. An attractive hypothesis is that AMNs in the lateral zebrafish thalamus serve, at least partially, as an LGN homolog, yet this hypothesis requires further testing.

### Mechanism of thalamic ODP

Visual input to a single eye imposes a turn bias in the contralateral direction during search behavior. Our data suggest that OFF-responses are primarily affected by ODP-like manipulation in the zebrafish thalamus, and the AMN OFF-responses are reduced in the hemisphere ipsilateral to the stimulated eye compared to the contralateral hemisphere (light-deprived eye). Interestingly, we also observe the opposite effect in greater thalamic neurons. We currently do not know the functional role of this subset of thalamic neurons in ODP. Yet, fewer OFF-responses are observed in greater thalamus relative to AMNs, suggesting a less influential role in ODP. We demonstrate activation of AMNs is sufficient to evoke motor responses suggesting a direct ability to impact motor performance.

Nonetheless, these divergent patterns of activity within the thalamus imply that both Hebbian (AMNs) and homeostatic (greater thalamus) plasticity mechanisms could be occurring, consistent with the multiple forms of plasticity occurring during mammalian ODP^75–77^. What remains lacking is whether these changes happen at retino-thalamic inputs, intrinsically within thalamic neurons, or in associated inhibitory circuits. Our data show increases in the number and strength of OFF response y279-positive AMNs in the hemisphere opposed to the turn bias direction and an opposite response profile in adjacent neurons in the greater thalamus, suggesting multiple forms of plasticity take place in the zebrafish thalamus. Regardless of how plasticity occurs, we propose a model where GABAergic AMNs regulate motor performance by a disinhibition model (Figure 7). Our previous work demonstrated that unilateral ablation of AMNs imposed a turn bias toward the intact hemisphere^26^, implicating that a contralateral projection needs to exist for our model to work. AMNs may inhibit contralaterally projecting premotor inhibitory neurons, which inhibitory glycinergic neurons have been described in the zebrafish hindbrain that regulate motor behavior and predominately inhibit contralateral neurons^78–80^. Alternatively, ipsilaterally projecting glycinergic neurons are also present in the zebrafish hindbrain, and future tracing experiments could demonstrate that AMNs project contralaterally, ultimately leading to the same behavioral effect.

### ODP regulation of search behavior

Previously, we demonstrated that the direction of motor asymmetry is not inherited, suggesting a role for environmental determinants^26,27^. Recently, we published a screen of etiologically relevant environmental factors showing that turn bias direction can be modulated by environmental experience. However, we did not elucidate a causative factor imposing left or right turn bias during search behavior^27^. Here we show that asymmetric visual experience instructs this outcome. Interestingly, the critical period interval is largely before swim bladder inflation when larvae are predominately on their side^81^. Therefore, larvae may naturally receive asymmetric visual input during this time, producing an environmental mechanism equivalent to the CP assay. As larvae would typically be randomly positioned in their native or laboratory environment, this model also explains the equal distribution of left versus right turners observed at the population level, providing a mechanism to generate behavioral randomness at the population level, which randomness and behavioral unpredictability has evolutionary advantages^82–84^. However, even following the CP assay, we note that a subset of individuals show motor asymmetry opposed to the expected turn bias direction based on positioning in the CP assay. This divergence could result from natural variation driving unique individualistic responses^85,86^. Alternatively, in our CP assay, the ‘occluded’ eye likely receives some visual input. In mammalian ODP experiments, an eye is typically surgically sutured to completely block visual experience. In our model, we benefit from avoiding surgical manipulations that could prompt inflammation and potentially interfere with plasticity^87^, yet this incomplete blockage may cause some behavioral noise in our experiments. Although a benefit to this approach is that future studies can address how changes in photo-stimulation during the CP affect plasticity, which would be challenging with surgical occlusion methods. Our work collectively establishes a powerful new vertebrate model to interrogate how visual experience modulates subcortical thalamic circuits and behavioral performance. Here we used this model to demonstrate neuron-level functional changes in the thalamus and implicate thalamic control of CP onset through inhibitory signaling.

## Methods

### Zebrafish husbandry

All experiments were approved by the West Virginia University Institutional Animal Care and Use Committee. Adult Tübingen long-fin (TL) zebrafish (*Danio rerio*) were used as a genetic background for all experiments. All experiments were performed within 14 days post fertilization (dpf) before sex determination. Larvae were raised at 28°C on a 14/10 hr light-dark cycle at 75 µW/cm^2^ intensity, in embryo media 3 (E3: 5mM NaCl, 0.17mM KCl, 0.33mM CaCl_2_, 0.33mM MgSO_4_-7H_2_O, and 1mM HEPES buffer pH 7.5) at a raising density of 40 larvae per 30mL E3 unless stated otherwise. Transgenic lines used were *Tg(y279: Gal4)*^26^, *Tg(UAS:Kaede)s1999t*^88^, *Tg(UAS:nsfB-mCherry)*^88^, *Tg(elavl3:H2B-GCaMP6f)*^41^, *Tg(elavl3:GCaMP6f)*^89^.

### Behavior tracking and analysis

#### Trajectory analysis

Larvae were recorded at 10 Hz using a µEye IDS1545LM-M CMOS camera, using infrared illumination (940nm, CMVision Supplies), and tracked using DAQtimer and custom software. Individual larvae were placed in 6-cm petri dishes and tracked over four simulation series of 30 seconds baseline illumination immediately followed by 30 seconds following the loss of illumination, with each recording set separated by 150s of untracked baseline illumination. We used three measures to establish motor behavior: Match Index, Total Turn Angle (TTA), and Bias Ratio (BR). *Match Index:* Proportion of trials that match the direction of the first dark trial bias. Individuals who have a rightward bias on the first trial and leftward bias on the subsequent trials would have a match index of 0, while individuals who have four trials of rightward bias would have a match index of 1. *TTA*: The sum of all angular trajectories regardless of direction. *BR*: The net turn angle is the sum of all angular trajectories with respect to direction (left turns being negative degrees change, while right turns are positive degrees) divided by the TTA. Therefore, individuals who only turn rightward will have a BR of 1, while those who only turn leftward will have a BR of -1. If an individual has an equal magnitude of leftward and rightward turns in a tracking period, their BR would be 0. Trials that had less than 10s of data were excluded from analyses. Individuals missing >2 baseline or >1 dark response(s) were excluded. Individuals missing the first dark trial were excluded from Match index calculations.

#### Critical period assay

For CP assay experiments, larvae were raised under standard conditions before, during, and after embedding. At 2 dpf larvae were manually dechorionated using forceps. Individuals were lightly anesthetized using dilute Tricaine (Sigma) until touch unresponsive and mounted in 1% low melting temperature agarose for all CP assay experiments. For embedding at later developmental stages (4-6 dpf) agarose was cut away from the mouth and gills to allow for adequate oxygenation and ion exchange. A 25% neutral density filter was placed under dishes to provide greater contrast between the eyes. For CP assay involving lack of visual stimulus, mounted individuals were kept in a blackout box. For the light-rotated visual stimulation CP assay, individuals were mounted on the inner lid of a petri dish with a neutral density filter placed on top of the dish and illuminated from below. Following removal from agarose, individuals were raised under normal conditions, and behavior was tested at 6-7 dpf for 2-4 dpf embedding and 9 dpf for 4-6 dpf embedding unless stated otherwise. To assess the level of asymmetric visual experience during the CP assay, individuals were laterally embedded in low melting temperature agar at 4dpf similar. We used a 455 nm LED (ThorLabs) at 10 µW/cm^2^ narrowed using a pinhole to a beam approximately the diameter of the larval eye. Using an optometer (International Light Technologies Model 2400 Optical meter), we measured light depression through the agar immediately adjacent to the larvae, the tail, and the eyes. Measures were standardized to the light intensity through the agar alone. For embedding larvae upright and providing a lateral light source. A black aryclic bar was glued to the center of a petri dish and larvae were agar embedded along the sides of the bar. LED arrays were symmetrically positioned on either side of the petri dish and experiment was carried out similar to typical CP assay experiments.

#### Extended timeline

Larvae were mounted and behavior tested as described above. Following behavior testing at 6-7 dpf, individuals were raised in our fish facility in system water. Larvae were raised in 300 ml of system water initially, with 200 ml added daily until 12 dpf where they were put on a slow drip water flow (∼1 drop/sec). Babies were fed 75 micron GEMMA until 12 dpf, when they transitioned to 150 micron GEMMA. Individuals were behavior tested at 14 dpf as described above, except using system water versus E3.

### Pharmacology

Neural activity was modulated using pentylenetetrazole (PTZ, Sigma, P6500), muscimol (Sigma, M1523), caffeine (Sigma, C0750), 4-aminopyridine (4-AP, Sigma, 275875), and gaboxadol (Sigma, T101). Stocks were prepared in molecular grade water. To test the impact on turn bias, larvae were treated with each drug diluted in E3 to working concentrations. The CP assay was conducted as previously mentioned. Drugs were replaced daily for all experiments. At 4 dpf, the drug(s) were removed, and fresh E3 was used for the rest of the experiment.

### Labeling experiments

#### Fluorescent in situ hybridization

Molecular Instruments HCR fluorescent in situ hybridization technology was used for all gene expression experiments. Company designed probe sets were generated based on provided accession numbers for gad1b (NM_194419; 17 probes), gad2 (NM_001017708; 20 probes), and vglut2b (NM_001009982, 17 probes). A company-validated kaede probe set (17 probes) was ordered to label AMNs in *Tg(y279:Gal4; UAS:Kaede)* larvae. At 3dpf, *Tg(y279:Gal4; UAS:Kaede)* larvae were fixed overnight at 4°C in 4% paraformaldehyde in 1x PBS and 0.1% Tween20. After fixation, embryos were washed in 1x PBS with 0.1% Tween 20, and labeling was conducted using HCR RNA-Fish protocol (Hageter et al. 2021). Imaging was performed on an Olympus Fluoview FV1000 confocal using a 40x oil immersion objective. To measure changes in brain activity, a fosab/c-fos (NM_205569, 27 probes) probe was designed by Molecular Instruments. Larvae were drug treated for 1 hour at 4dpf. Following treatment, larvae were fixed and labeled for fosab. Images were captured on a ThorLabs Confocal with a TOPTICA iChrome CLE 50 laser and 20x objective using consistent laser power and gain settings across all samples. A consistent plane in the brain was captured for all larvae. For analysis, the whole brain area mean intensity was quantified using ImageJ and standardized to control larvae.

### Functional Imaging

#### Calcium imaging

Calcium imaging was performed across whole single planes, using *Tg(y279: Gal4); Tg(UAS:nsfB-mCherry); Tg(elavl3:H2B-GCaMP6f)* or *Tg(elavl3:GCaMP6f)* for nuclear-localized or regional calcium recording, respectively. Larvae were raised in 200µM PTU from 1 dpf until 24 hours before imaging. Behavior was run to confirm turn bias direction at 6 dpf. For enucleation experiments, larvae were anesthetized and unilaterally enucleated at 5 dpf in 1x Evans and returned to E3 approximately 24 hours post-enucleation. For nuclear-localized calcium imaging, larvae were mounted in the CP assay. At 7 dpf, larvae were lightly anesthetized using tricaine and mounted in 2% low-melting temperature agarose. Experiments were performed using a Scientifica Vivoscope two-photon, a 16x water immersion objective, and Spectra-Physics MaiTai laser tuned to 980nm. Visual stimulation was provided using a Thorlabs mounted 660-nm LED at 70 µW/cm^2^. For each larva, a 1-minute light ON, 1-minute light OFF, and 1-minute light ON stimulus series was repeated twice per larvae with a 3-minute red light ON interval between replicates. Stimulation series were preceded by 5 minutes of constant red light illumination to adapt larvae. We captured a single plane of the thalamus using *Tg(y279: Gal 4; UAS: nsfB-mCherry)* as a guide for AMNs and surrounding brain regions. We chose a single plane based on neuroanatomic structure for imaging the optic tectum. Image acquisition was performed using ScanImage software at ∼1.01 Hz. All images were registered using the TemplateMatching plugin for ImageJ^90^. For nuclear-localized calcium imaging, we analyzed fluorescence intensity changes by manually drawing an outline of each cell and extracting mean gray values from those ROIs using a sum projection and the Romano-et-al toolbox in Matlab. A custom program was written in Matlab to calculate (*F*_t_ − *F*_0_)/*F*, (∆*F*/*F*), where F_0_ was the mean gray value during the baseline period. Neurons with an ∆*F*/*F* greater than 3 standard deviations of the baseline for 3 successive frames following complete light extinction were classified as OFF-responses, following the return of illumination, ON-responses, and both OFF/ON-Responses. Cells were categorized into regions based on localization and using Mapzebrain regions^43^. Using a composite image of GCaMP6f and the y279 reporter, cells were classified to either be y279 positive or negative. Analysis for regional recordings was performed by outlining ROIs in ImageJ and using the “plot z-axis profile” function to extract gray values and calculate ∆*F*/*F* in Excel. For mosaic calcium imaging, embryos at the single cell stage were injected with 20 pg of UAS:NLS-GCaMP6s^26^ plasmid and co-injected with 25pg of tol1 mRNA. Larvae were raised, behavior tested, imaged, and analyzed as above.

***iGluSnfer*** Single-cell stage *Tg(y279: Gal4; UAS:nsfB-mCherry)* embryos were co-injected with 20 pg of UAS: 14xUAS-iGluSnf^91^ and 25 pg of tol1 mRNA. Larvae were raised in 200µM PTU from 1 dpf -5dpf and enucleated at 5 dpf. Larvae were mounted and imaged, similar to the calcium imaging experiments. Images were captured at 2.00 Hz. We calculated ∆*F*/*F* from a region lateral to the y279 reporter in the adjacent neuropil and a posterior position as a control.

### Optogenetics

Single-cell-stage *Tg(y279:Gal4; UAS:Kaede)* embryos were co-injected with 25 pg of UAS:TagRFP-v2a-paGFP (kind gift from Harold Burgess lab), UAS:ChEF-v2a-mCherry^63^, or UAS:GtACR2-tdTomato plasmid^64^ (Addgene plasmid #124236) and 25 pg of tol1 RNA. Individuals were raised in 200 μM PTU until 7dpf. At 7dpf, larvae were mounted in 2% low-melt agarose with the tail cut free and imaged using a Scientifica two-photon Vivoscope with Spectra-Physics MaiTai laser and 16x water immersion objective. Image acquisition was performed with ScanImage software through MatLab. The laser was tuned to 940 nm (15 μW) and was focused on a single plane through the AMNs at 10x zoom. Each larva went through a stimulation series that consisted of 3 sequential recordings: 5 minutes of darkness followed by either a 20-second recording of dark (no laser), 940nm, and 1040nm stimulation, respectively. A µEye IDS1545LM-M CMOS camera recorded tail movement with a 12 mm lens, a long-pass 780 nm filter, and a short-pass 900 nm filter (Edmund Optics), and infrared illumination was provided using a Thorlabs fiber optic 850nm LED driver and fiber optic cable (M28L02). The camera’s field of view was set to record the tail of each larva, and DAQtimer software was used to conduct real-time tracking of tail movement at 10 fps. Points were manually assigned to the tip of the tail for each recording frame as a measure of tail displacement from the original tail position.

#### Cell death analysis

To determine if the optogenetic stimulation resulted in cell damage, *Tg(y269:Gal4; UAS:mCherry)* embryos were raised in 200 μM PTU until 7dpf and mounted as described above. Control individuals were put through the optogenetic stimulation series. To induce photo-damage as a control, the 2-photon laser was tuned to 940nm (80 μW) and zoomed in 70x between the AMNs to focus laser power during a 10-second scan. Cell death was detected using acridine orange (AO) for 20 minutes (5 ng/μL in E3), followed by multiple washes in E3 for an hour. Finally, larvae were mounted in 2% low-melt agarose and imaged using a Thorlabs Confocal (TOPTICA iChrome CLE 50 lase, 10x objective).

## Statistical Analysis

Analysis was performed in Matlab, RStudio, and Graphpad. Data in the figures and text represent means +-SEM. All t-tests were two-sided. Wilcoxon Signed Rank Sum tests were used for the match index compared to 0.5. Normality was determined using the Shapiro-Wilks test. Multiple comparisons were correcting using a Bonferroni correction. Chi-square statistics were performed in Graphpad.

## Acknowledgments

We want to thank Sarah Ackerman and Andrew Dacks for their helpful comments on the manuscript. We thank Johnathan Huff for assistance with zebrafish care and preparation for CP assays. In addition, we thank Misha Ahrens, Harold Burgess, Steve Finkbeiner, and Jeremy Linsley for resources, including fish lines and plasmids.

## Author contributions

EH, JH, and JS conceived the project and wrote the manuscript. JH and JS performed experiments. All authors approve the submitted version.

