## Supplemental material for "Thalamic regulation of a visual critical period and motor behavior"

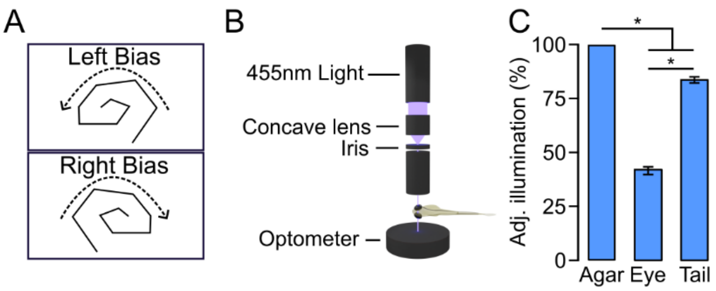


Supplemental Figure 1

**Supplemental Figure 1: (A)** Representative left and right biased trajectories following the loss of light. **(B)** Schematic of setup used to test zebrafish light permeability. **(C)** Normalized percent light transmission for the tail (N=10) and eye (N=10) relative to low melting temperature agar. Asterisk $p<0.05$ two-tailed t-test.

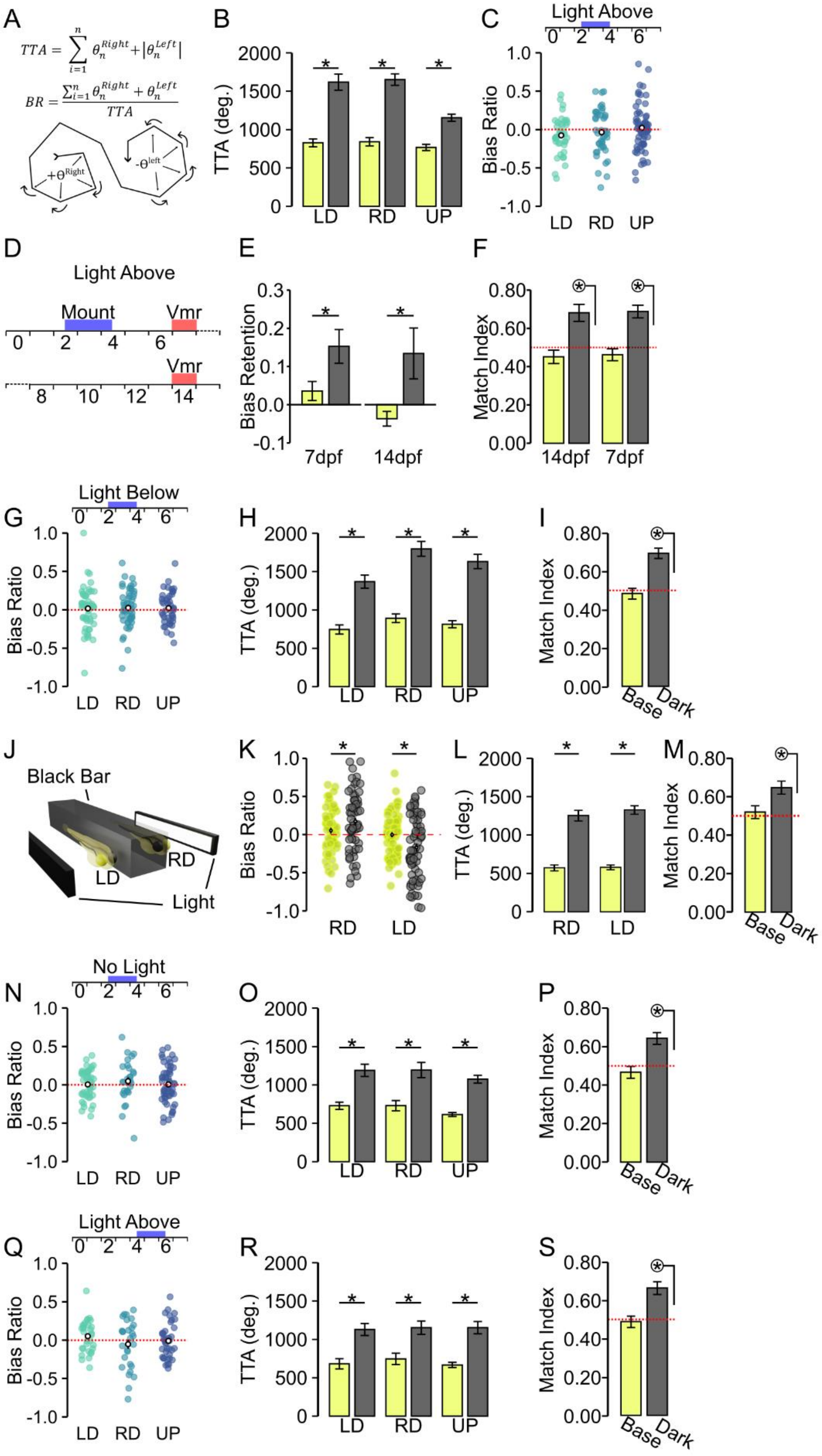


Supplemental Figure 2

**Supplemental Figure 2: Zebrafish critical period is visually mediated and developmentally restricted. (A)** Total turn angle (TTA) and bias ratio (BR) calculation. CP assay from 2-4 dpf, light overhead, showing baseline total turning angle (TTA) **(B)** and bias ratios **(C)**, representing Figure 1H (LD=39, RD=40, UP=62). Asterisk $p < 0.05$ two-tailed t-test. **(D)** Timeline for extended testing of motor asymmetry. **(E)** Bias retention (average BR with all recording from left individuals multiplied by '-1' to standard turn direction across groups) according to mounted orientation for baseline and dark responses. Asterisk $p < 0.05$. **(F)** Match index for experiment outlined in D-E. Circled asterisk indicates $p < 0.05$ one-sample Wilcoxon signed rank test to 0.5 showing turn bias direction preserved following the CP assay up to two weeks post-fertilization. **G-I)** Baseline values for 2-4 dpf CP assay, light below showing average bias ratio **(G)**, total turning **(H)**, and match index **(I)**. (LD N=51, RD N=53, UP N=42). **(J)** Representative diagram for upright CP assay using a black spacer to produce visual asymmetry. **(K-M)** Motor metrics from upright CP assay showing bias ratio **(K)**, total turning **(L)**, and match index **(M)**. Baseline recordings from 2-4 dpf CP assay with no photic stimulation (LD N=44, RD N=28, UP N=54) showing **(N)** Baseline BR, **(O)** TTA, and **(P)** match index. **(Q-S)** Same as above for 4-6 dpf CP assay experiment (LD N=31, RD N=28, UP N=39). Asterisk $p < 0.05$ two-tailed t-test. Circled asterisk one-sample Wilcoxon signed rank test to 0.5.

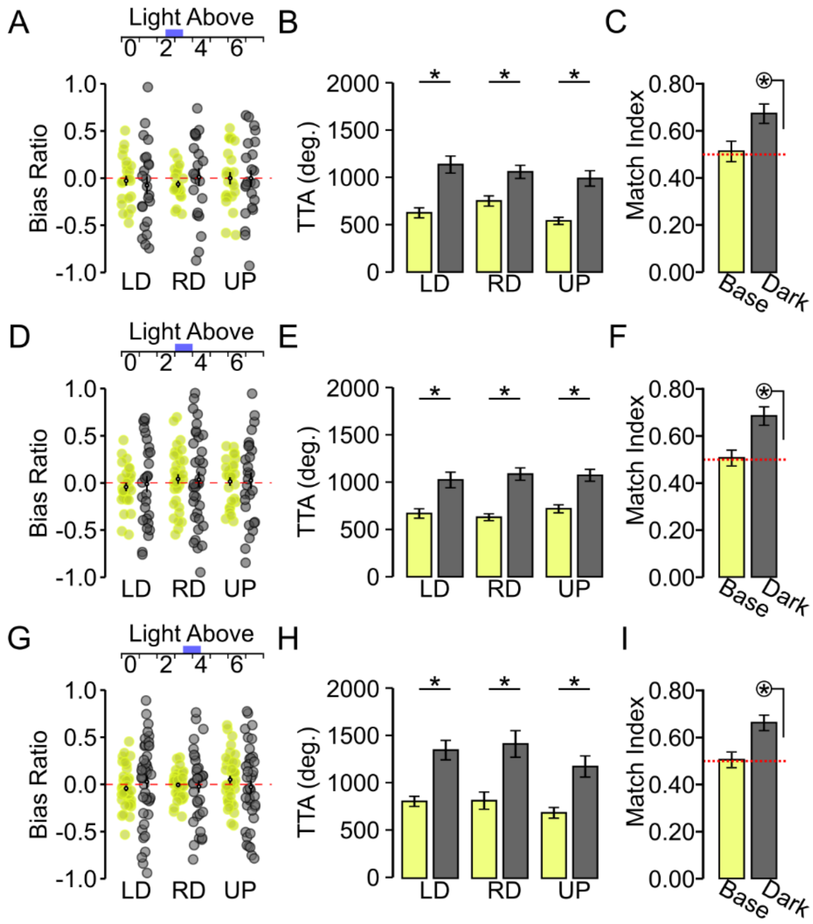


Supplemental Figure 3

**Supplemental Figure 3: Shorter periods of asymmetric visual control do not shift turn bias. (A)** BR, **(B)** TTA, and **(C)** match index (N=53, groups combined) for baseline (yellow) and dark (gray) responses for CP assay from 2-3 dpf (LD N=23, RD N=22, UP N=23). **(D)** BR, **(E)** TTA, and **(F)** match index (N=79, groups combined) for baseline (yellow) and dark (gray) responses for CP assay for 3 dpf (LD N=27, RD N=35, UP N=27). **(G)** BR, **(H)** TTA, and **(I)** match index (N=95, groups combined) for baseline (yellow) and dark (gray) responses for CP assay from 3-4 dpf (LD N=43, RD N=35, UP N=38). Asterisk indicates $p<0.05$ two-tailed t-test. Circled * indicates $p<0.05$ one sample Wilcoxon signed rank test to 0.5.

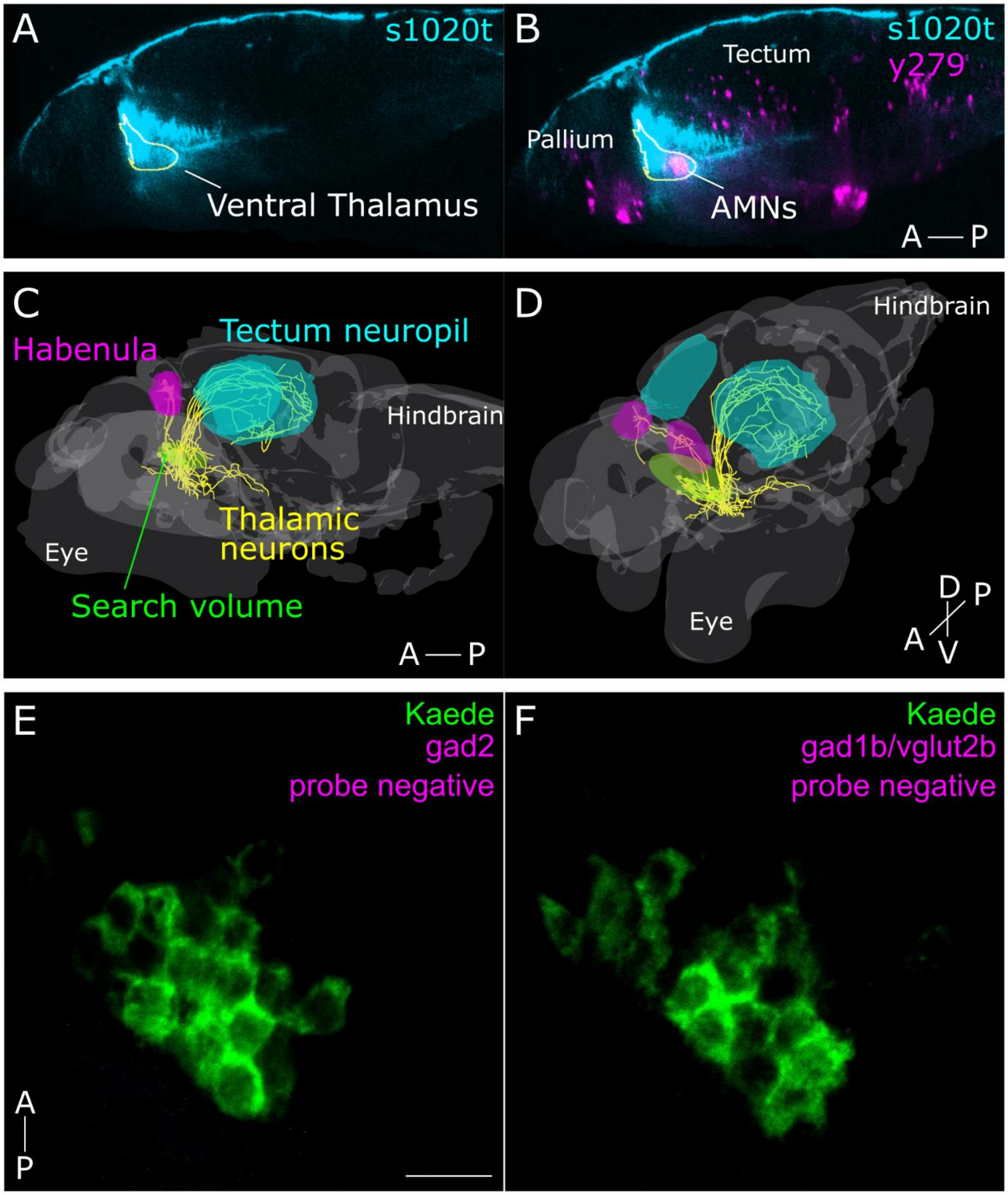


Supplemental Figure 4

**Supplemental Figure 4: AMNs reside in the thalamus**. **(A-D)** Images constructed from the mapzebrain atlas for zebrafish neuroanatomical markers and single neuron traces. **(A-B)** Colocalization of thalamic marker s1020t (cyan) and y279 (magenta). Outline is the mapzebrain segmented region for the ventral thalamus. Atlas comparison is optimal as both lines are Gal4 drivers and are not compatible for co-localization in vivo. **(C-D)** 3D reconstruction of single neuron traces showing neurons identified with soma residing in the AMN region (green search volume) that project to the habenula (magenta, N=2 identified neurons) or the tectum neuropil (cyan, N=10 identified neurons). All identified neurons mirrored to the left hemisphere for ease of visualization. **(E-F)** Negative control labeling for custom GABAergic and glutamatergic HCR FISH probes. *Tg(y279:Gal4; UAS:kaede)* larvae were labeled for kaede mRNA (green) as normal (probe and hairpin) and single plane confocal scans captured showing a single hemisphere AMNs. To test custom HCR probes either b1 (N=4 larvae) (F) or b4 (N=4 larvae) (G) hairpins were used with no probe, respectively (magenta). Fluorophore tagged to hairpins in HCR technology. Scale bar 10 µm.

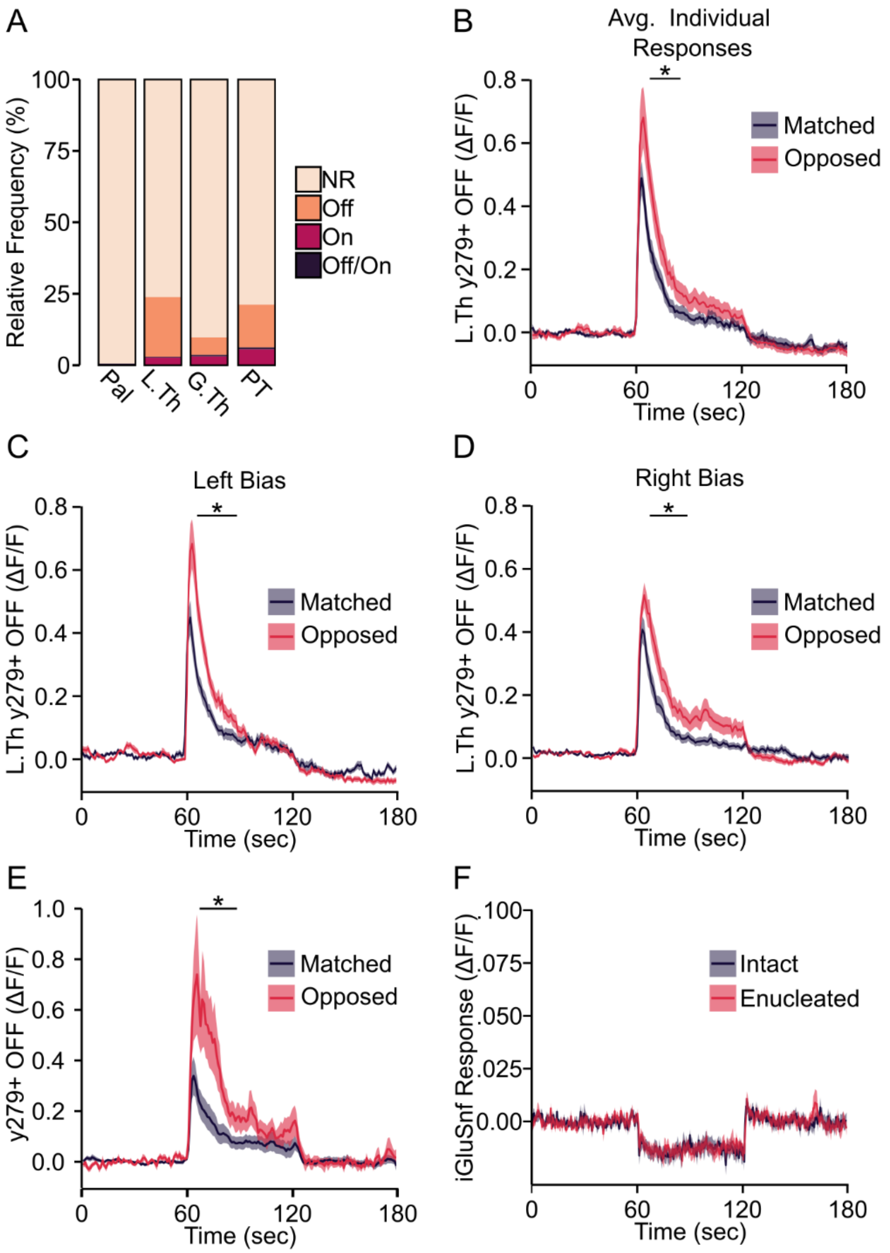


Supplemental Figure 5

**Supplemental Figure 5: Regionalized OFF-Responses coincide with turn bias. (A)** Response type distribution for all regions (tan=No Response, orange = OFF response, red=ON response, Purple = OFF/ON response). **(B-D)** Differential analysis of lateral thalamus OFF-response calcium imaging data. **(B)**. All single neuron OFF-responses were averaged per recorded individual (N=17) for comparison between matched and opposed hemispheres. **(C-D)** Photo-mediated calcium response for left-biased (Matched N=45, Opposed, N=55) **(C)** and right-biased (Matched N=30, Opposed, N=49) **(D)** individuals. **(E)** Responses from mosaic analysis using *UAS:BGi-nls-GCaMP6s* constructed injected into *Tg(y279:Gal4)* larvae (Matched N=17 neurons; Opposed N=19 neurons; recorded from N=11 left bias and N=10 right biased larvae). **(F)** iGluSnfer responses from intact (red) and enucleated (grey) hemispheres recorded from neuropil 50 µm posterior to the AMNs (N=9). Quantification from same individuals used in Figure 4F. Line shows average standardized fluorescence and envelope ±SEM. Asterisk indicates $p<0.05$ for ten consecutive seconds, two-tailed t-test.

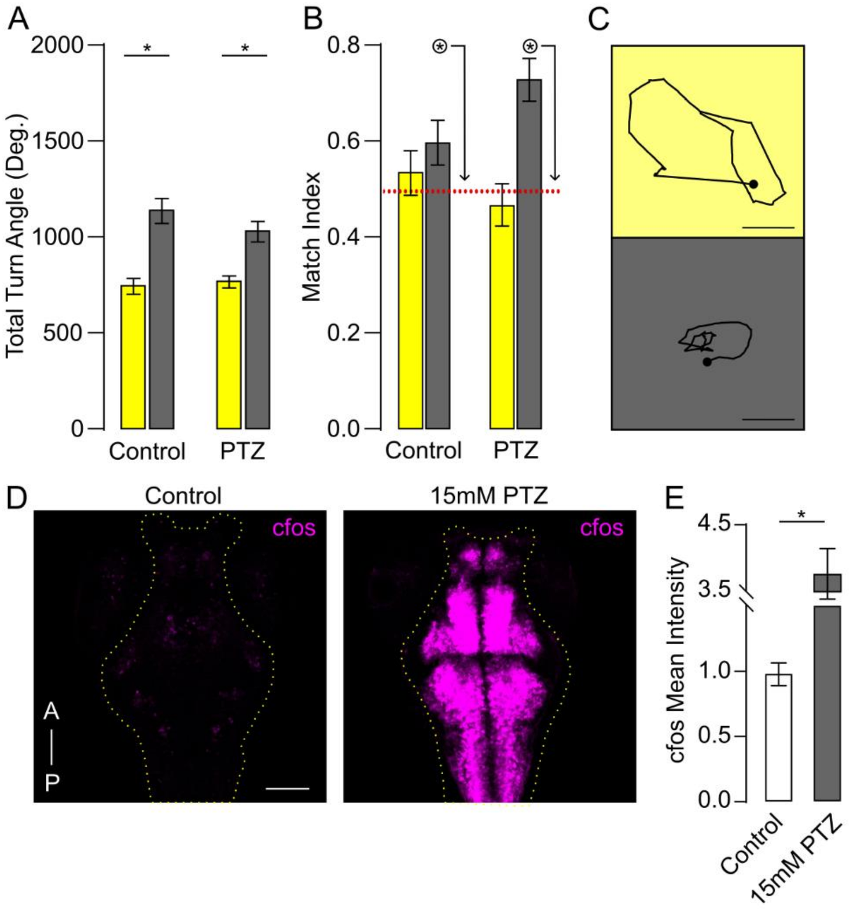


Supplementary Figure 6

**Supplemental Figure 6: GABAergic signaling does not influence circling behavior.** Average total turning **(A)** and MI **(B)** of individuals in baseline (yellow) and dark (gray) for control (N=41) and treated with PTZ (N=40) from 2-4 dpf CP assay. **(C)** Representative behavioral trace of an individual treated with PTZ from 2-4 dpf in baseline (yellow) and dark (gray). Scale bar 10mm. **(D)** Representative c-fos HCR FISH labeling following high-dose PTZ treatment compared to untreated control. Single plane confocal scans of 4 dpf larvae. Yellow outline shows the area analyzed. Scale bar 100 µM. **(E)** Quantification of D. Untreated controls (N=10) and 15mM PTZ (N=10). Values standardized to control. Asterisk with the line shows $p<0.05$ between groups. Circled Asterisk $p<0.05$ to 0.5.

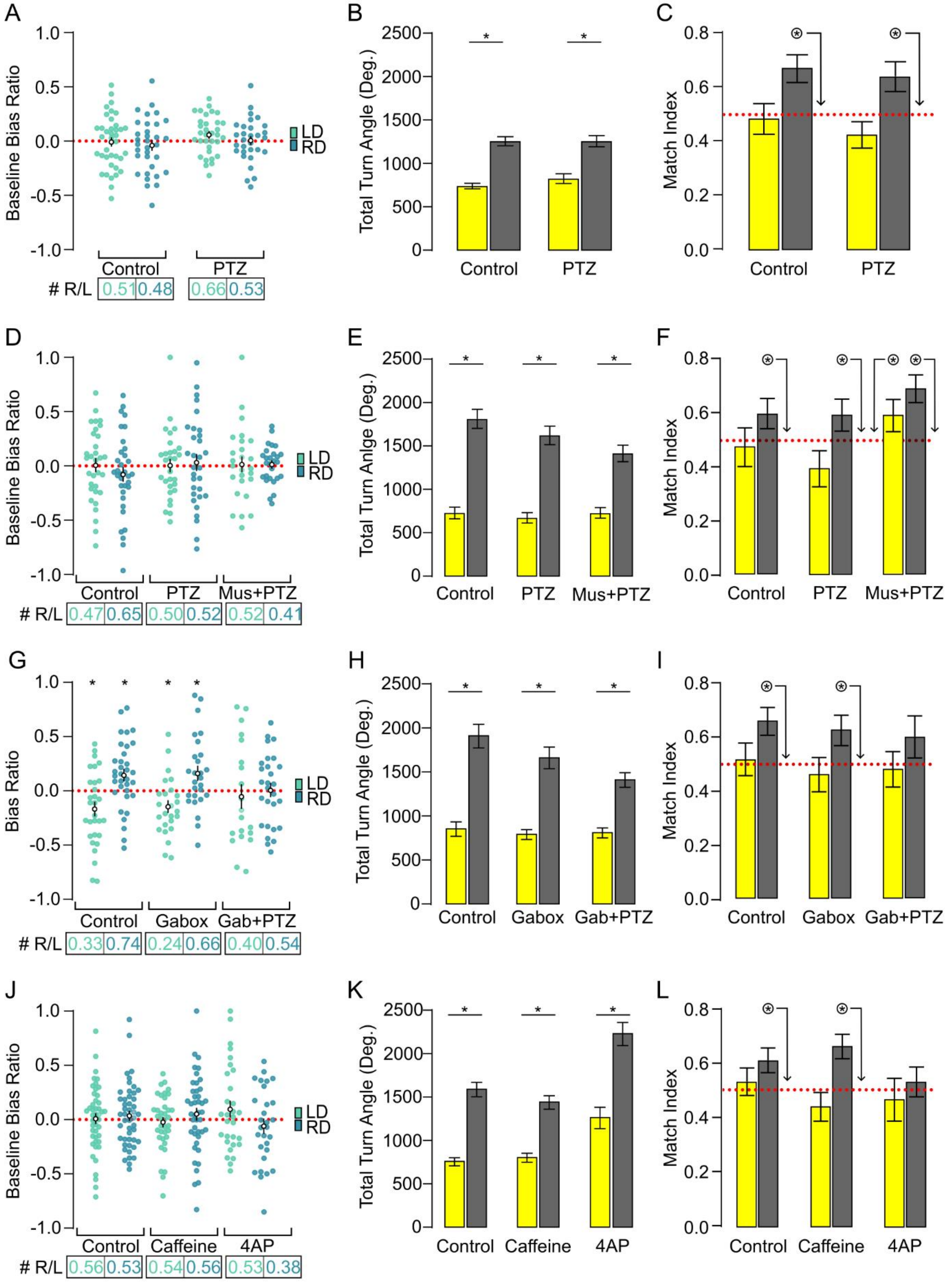


Supplementary Figure 7

**Supplemental Figure 7: GABAergic modulation of turn bias plasticity.** Motor parameters for drug treatment experiments showing average BR (**A,D,G,J;** bottom numbers show the R/L ratio of right to left biased individuals), total turning **(B,E,H,K),** and match index **(C,F,I,L)**. RD and LD groups combined for TTA and MI analysis and shows baseline (yellow) and dark response (grey) data. **(A-C)** Baseline responses for PTZ treatment (control RD N=33, LD N=39; PTZ RD N=30, LD N=33). **(D-F)** Baseline responses for PTZ and muscimol cotreatment (control RD N=35, LD N=32; PTZ RD N=31, LD N=30; Muscimol+PTZ RD N=27, LD N=26). **(G-I)** Gaboxadol treatment during the CP assay. **(G)** Average dark response BR for control (RD N=34, LD N=30), Gaboxadol (RD N=27, LD N=25), and Gaboxadol+PTZ (RD N=28, LD N=20). Average total turning **(H)** and MI **(I)** (control N=64, Gaboxadol N=52, and Gaboxadol+PTZ N=48). **(J-L)** Baseline responses for other activity-modulating drugs (control RD N=47, LD N=46; caffeine RD N=43, LD N=37; 4-AP RD N=29, LD N=30). Asterisk with the line shows $p<0.05$ between groups. Circled Asterisk $p<0.05$ to 0.5.

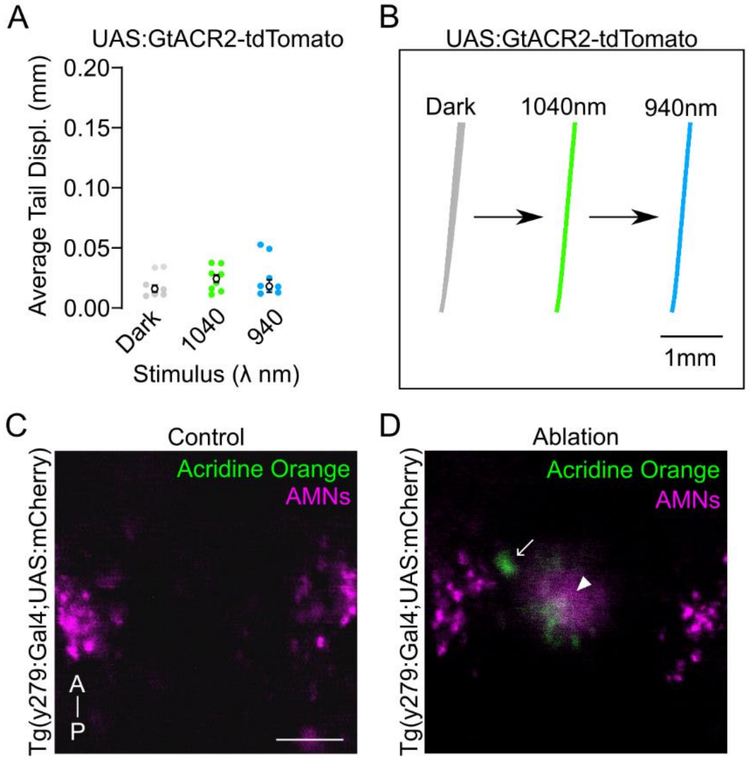


Supplemental Figure 8

**Supplemental Figure 8: Optogenetic characterization of AMNs. (A-B)** Optogenetic inhibition of AMNs using GtACR2 showing tail displacement **(A)** and illustrative tail trace examples **(B)** (N=8). Test for cell death following two-photon stimulation **(C-D)**. Images are single-plane confocal scans of AMNs following either parameter used for optogenetic stimulation experiments **(C)** or high-power laser exposure as a control for cell damage **(D)**. Larvae labeled with acridine orange (green). Arrow indicates acridine orange-stained cells. Arrowhead shows an ablation scar. Similar patterns were observed in control (N=4) and ablation (N=4) experiments. Scale bar 20 μm.

| OFF-Responses | | | | |
|---|---|---|---|---|
| Region | match N | oppose N | $X^2$ | $p$ |
| L.Th y279+ | 75 | 104 | 4.698 | 0.0302 |
| L.Th y279- | 28 | 38 | 1.515 | 0.2184 |
| G.Th | 61 | 79 | 2.314 | 0.1282 |
| PT | 167 | 143 | 1.858 | 0.1728 |

| ON-Responses | | | | |
|---|---|---|---|---|
| Region | match N | oppose N | $X^2$ | $p$ |
| L.Th y279+ | 10 | 15 | 1 | 0.3173 |
| L.Th y279- | 2 | 4 | 0.667 | 0.4142 |
| G.Th | 41 | 29 | 2.057 | 0.1515 |
| PT | 50 | 65 | 1.957 | 0.1619 |

**Supplemental Table 1:** Distribution of OFF and ON-responsive neurons between matched and opposed hemispheres. Chi-square analysis performed based on observed neuron distributions compared to an expected equal distribution based on total counts per region.
